# Metal homeostasis is remodeled in response to different quiescence triggers

**DOI:** 10.64898/2026.05.21.727032

**Authors:** Ananya Rakshit, Samuel E Holtzen, Allegra T. Aron, Aidan T. Pezacki, Smitaroopa Kahali, Sujit K. Das, Lynn Sanford, Martina Ralle, Ankona Datta, Amy E Palmer

**Affiliations:** Department of Biochemistry, University of Colorado Boulder, Boulder, CO, 80303, USA; BioFrontiers Institute, University of Colorado Boulder, Boulder, CO, 80303, USA; Department of Molecular Cellular and Developmental Biology, University of Colorado, Boulder, CO, 80309; Department of Chemistry, University of California, Berkeley, Berkeley, CA 94720, USA; Department of Chemistry, Princeton University, Princeton, NJ, 08544, USA; Department of Chemical Sciences, Tata Institute of Fundamental Research, 1 Homi Bhabha Road, Colaba, Mumbai, 400005, India; Department of Molecular and Medical Genetics, Oregon Health & Science University, Portland, OR, 97239, USA

**Keywords:** metal homeostasis, quiescence, zinc deficiency, RNA seq, labile metal pool

## Abstract

Cells can enter a reversible non-proliferative state called quiescence either spontaneously or in response to nutrient deprivations. Metal ions are essential nutrients and play wide-ranging regulatory and signaling roles in biological processes. We previously showed that zinc is an essential nutrient for the mammalian cell cycle as zinc deprivation drives cells into quiescence, and this quiescent state is associated with changes in iron, copper, and manganese, suggesting broad remodeling of metal homeostasis. Here we examine whether metal remodeling is a general feature of quiescence by inducing quiescence via different triggers (zinc deficiency, serum starvation, and growth factor withdrawal) in MCF10A cells. Fluorescence microscopy and elemental analysis reveal significant trigger-dependent changes in the labile and total metal pools of quiescent cells. To gain insight into these differences, we carried out RNA sequencing and differential expression analysis, focusing on metal associated, metal regulatory, and metal homeostasis genes. While core quiescence pathways are shared across triggers, quiescence states remain molecularly distinct. A significant percent of 2458 metal homeostasis annotated genes are differentially expressed including 55% in starvation-induced quiescence, 50% in zinc deficiency-induced quiescence and 21% in growth factor withdrawal-induced quiescence. Our results also showed unique alteration of genes involved in major metal dependent processes including antioxidant activity, oxidative phosphorylation, heme metabolism, and chromatin accessibility in different quiescence states. Overall, this work demonstrates that metal homeostasis is systematically rewired during cellular quiescence with associated effects on genes that regulate critical biological processes.

**Significance Statement:** Cells constantly evaluate their nutrient and energy status and integrate these signals into proliferation-quiescence decisions. Quiescence prevents cells from passing damage to daughter cells. Metals are essential micronutrients for biological processes. While limitation of zinc can drive cells into quiescence and alter other metals, how metals are remodeled and whether this is a common feature of quiescence was unknown. Here we examined three quiescence triggers by modifying growth media and serum, the primary source of metals. We report significant changes in labile and total Cu, Fe, Zn and Mn pools, and metal associated genes in response to distinct quiescence triggers. These changes converge on mitochondrial function, cellular antioxidant activity, and heme biosynthesis in a trigger specific manner.

## Introduction

The eukaryotic cell cycle is a highly regulated process that is largely conserved from yeast to mammals.(1) The cell cycle involves controlled progression from a gap phase (G1) to DNA replication (S) to a second gap phase (G2) and finally mitosis (M), resulting in two daughter cells. A well-recognized feature of the cell cycle is the capacity of cells to enter quiescence, a reversible non-proliferative state (also known as G0) after mitosis. A small population of cells can enter quiescence spontaneously,(2) but quiescence can also be induced by a variety of factors, including growth factor withdrawal, serum starvation, contact inhibition, and restriction of glucose or amino acids.(1, 3, 4) We recently demonstrated that quiescence is also triggered by zinc (Zn) deficiency.(5) Zn deficient quiescent cells also showed changes in other essential metal ions, such as iron (Fe), copper (Cu), and manganese (Mn), suggesting broad changes in metal homeostasis. Whether remodeling of metal ions is a general feature of quiescence has not been explored.

Although quiescence is often associated with dormancy, it is well established that quiescence is actively maintained and involves reprogramming of transcription, signaling, and metabolism.(2, 4, 6, 7) Numerous studies suggest quiescence is not a single state but rather a collection of heterogeneous states. Fibroblast cells induced into quiescence by different triggers (serum starvation, adhesion inhibition, contact inhibition) exhibit distinct transcriptional signatures, suggesting trigger-specific transcriptional programs.(4) This observation has been reinforced by subsequent studies looking at a broader spectrum of triggers.(2) Furthermore, quiescence is characterized by varying degrees of depth, where cells deeper in quiescence require stronger stimulation and/or more time to reenter the cell cycle.(8) Despite the differences, there are also gene expression changes shared by quiescent states suggesting that there is a common set of core changes.(2, 4)

Transition metals such as Zn, Fe, Cu, and Mn are essential micronutrients that are required for mammalian cells to grow and thrive. Beyond their structural and catalytic roles, Zn, Fe, Mn and Cu have been increasingly recognized as signaling ions and have the potential to influence transcriptional regulation.(9–13) Given that limitation of other nutrients such as glucose and amino acids induces quiescence and transcriptional reprogramming,(1, 14) it is logical that limitation of metals could induce quiescence. However because quiescence is associated with catabolism,(1, 15, 16) potential mobilization of metal ions from proteins or internal stores is possible. We previously showed that withdrawal of Zn led to quiescence in mammary epithelial cells (MCF10A) and that resupply led to synchronous cell cycle re-entry.(5) Intriguingly, the Zn deficiency-induced quiescent cells were characterized by changes in the total cellular content of Fe, Cu, and Mn, suggesting broad remodeling of metal homeostasis. This observation led to our overarching research question of whether reprogramming of metal homeostasis is a common feature of quiescent cells and whether different triggers might lead to different metal remodeling signatures.

In this study we examine whether cellular quiescence is associated with changes in the total and labile metal pool. We discovered significant perturbation of Zn, Fe, Cu and Mn, firmly establishing that metal content is altered in quiescence. Importantly, distinct quiescence triggers induced unique metal profiles, indicating that the nature of the trigger shapes metal homeostasis. To understand these differences, we analyzed the transcriptional signature of quiescent cells, focusing on changes in metal homeostasis and metal regulatory genes. We observed widespread transcriptional remodeling with starvation showing the highest number of differentially expressed genes compared to spontaneous quiescence, followed by Zn deficiency, followed by growth factor withdrawal. All three quiescence triggers induced broad changes in metal associated genes, where differential regulation of metal homeostasis genes suggest functional differences in key biological processes, including energy metabolism, oxidative stress, chromatin accessibility, and iron metabolism. Overall, this study demonstrates that cellular quiescence is associated with remodeling of the metallome and metal associated genes in a trigger-specific manner.

## Results

### Zn deficiency-induced quiescence is associated with large transcriptional changes

Metals are essential nutrients that are supplied to cells in serum and media. We previously showed that removal of Zn with the chelator tris(2-pyridylmethyl)amine (TPA), without perturbing other nutrients was sufficient to induce quiescence.(5) But how gene expression is remodeled in this Zn deficient quiescent state and how this state compares to other quiescent states is unknown. To answer these questions, we carried out next generation RNA sequencing on different populations of quiescent cells, including spontaneous quiescence (triggerless) and cells driven into quiescence by three different triggers, Zn deficiency (ZD), serum starvation (Starv), and growth factor (GF) withdrawal. ZD involved treatment with TPA (3 µM), Starv involved removal of serum, insulin and growth factors, while GF involved selective removal of growth factors and insulin. We first evaluated the optimal treatment time for obtaining the highest percentage of quiescent cells. To accomplish this, MCF10A cells expressing p21-mCitrine and geminin-mCherry were treated with quiescence triggers for 24h, 48h and 72h. Cells with high p21-mCitrine and low geminin-mCherry were identified as quiescent, as established previously.(2) The 48h treatment yielded the highest percentage of cells in the quiescent population (***SI Appendix*, Fig. S1**), with ZD, GF and Starv giving rise to 82%, 45% and 55% quiescent populations, respectively. 17-25% of cells grown in minimal media went quiescent spontaneously, as observed previously.(2, 17)

To characterize the transcriptome of different quiescent cell populations, cells were FACS sorted for high p21-mCitrine and low geminin-mCherry, thus enriching the population of quiescent cells (**Fig. 1*A**, SI Appendix*, Fig. S1**), and subjected to RNA sequencing and differential gene expression analysis. Principal component analysis revealed distinct populations of cells that clustered primarily according to quiescence condition, with GF, Spont, and Cycling displaying the least intergroup variation compared to ZD and Starv (**Fig. 1*B***). Consistent with such clustering, ZD and Starv conditions give rise to greater transcriptional changes as reflected in broader distributions of log2Foldchange (Log2FC) values for gene expression relative to Spont (**Fig. 1*C***). Further investigation of gene expression upon induction of quiescence revealed distinct transcriptional signatures for different triggers, and interestingly, half or more of the transcriptional changes in Starv and ZD conditions are unique to those triggers (5687 and 4392 differentially expressed genes (DEGs), respectively) (**Fig. 1*D-E**, SI Appendix*, Dataset S1**). Given that serum starvation involves removal of growth factors and essential micronutrients like Zn, Fe, and Cu, such widespread transcriptional changes were expected, but such drastic changes induced by ZD was surprising given the specificity of the Zn chelation trigger. Despite the different signatures, there were 637 upregulated and 834 downregulated DEGs that were common among all quiescence triggers. Kyoto Encyclopedia of Genes and Genomes (KEGG) pathway analysis of common genes indicated shared downregulation of cell cycle genes and RNA processes (splicing, degradation, ribosome biogenesis) and upregulation of lysosomal and autophagic processes (**Fig. 1*F***), as reported previously for quiescence.(2, 4, 6–8, 18, 19) Thus each of the triggers yielded a common quiescence transcriptional profile amid other distinct widespread transcriptional changes.

**Figure 1.**
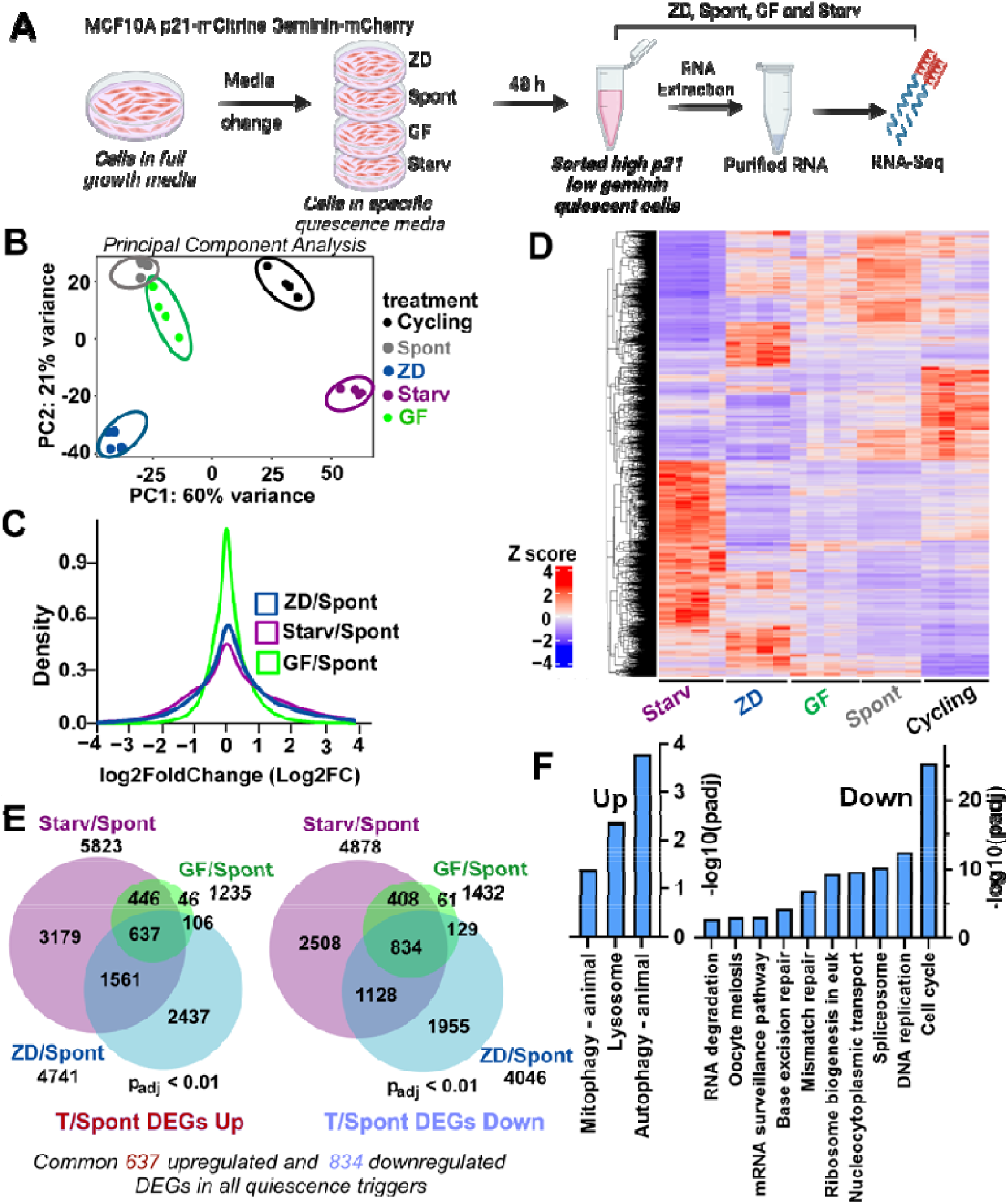
Analysis of differential gene expression for different quiescence subtypes. (A) Schematic overview of the bulk RNA-sequencing experiment performed on MCF10A p21-mCitrine geminin-mCherry cells grown under different media conditions. ‘Spont’ refers to sorted spontaneously quiescent cells grown in Minimal media (MM), ZD (MM + 3 µM TPA), GF (MM lacking growth factor and insulin); Starv (MM lacking serum, growth factor, and insulin). ‘Cycling’ refers to unsorted cycling cells grown in MM. N=4 biological replicates per condition. (B) Principal component analysis of RNA-Seq results from all samples. (C) Density plot showing log2Foldchange (Log2FC) distributions of differentially expressed genes ranging from -4 to 4 across all pairwise comparisons of quiescence trigger vs spontaneous quiescence (T/Spont). (D) Heatmap showing expression profile of ∼ 8000 genes regulated during quiescence across all RNA-Seq samples. Relative gene expression levels are shown as Z-scores and color-coded from violet (low expression) to red (high expression), ranging from -4 to 4, with white indicating zero. Each row corresponds to a gene and each column to a biological replicate. (E) Venn diagrams depicting upregulated and downregulated differentially expressed genes (p_adj_< 0.01) from pairwise comparisons of all T/Spont. (F) KEGG pathway analysis of 637 commonly upregulated and 834 commonly downregulated differentially expressed genes across all quiescence, plotted on a –log10(padj) scale.

Comparison of quiescence triggers to cycling cells (T/Cycling) also revealed changes in gene expression, including both trigger specific and shared genes **(*SI Appendix*, Fig. S2A-B**). RNA sequencing of serum starvation and cycling samples was performed separately from the remaining samples and DESeq2 analysis was performed without batch correction (see Methods, ***SI Appendix)***. Despite being processed differently, KEGG analysis of 1,230 shared downregulated DEGs in the T/Cycling comparison identified significant downregulation of ‘Cell cycle’ and four additional pathways consistent with previous reports **(*SI Appendix*, Fig. S2C**).(2, 4) We also compared our Starv/Spont data with a reanalyzed dataset from Min et al (2) and observed strong agreement, with 73% (4132/5660 genes) overlap between their reported DEGs and those identified in our Starv/Spont analysis (see Methods and ***SI Appendix*, Fig. S2D**). Overall, our analysis confirms that similar to other quiescent states, zinc deficiency is associated with significant changes in gene expression, and the profile is quite different from other quiescent states.

### Quiescence alters the total and labile metal pools

In mammalian cells many transition metals are buffered such that they exist in a pool where they are tightly bound to proteins and other biomolecules and as weekly-bound labile pool, which collectively constitute the intracellular total metal pool.(20, 21) We previously found that ZD quiescent cells (24 h) had altered levels of total Fe, Cu, and Mn as measured by Inductively Coupled Plasma Mass Spectrometry (ICP-MS) compared to cycling cells.(5) To evaluate whether different quiescence triggers induce different changes in the total metal status of cells, we treated cells for 48 hrs, then digested the cells and measured total metal content by ICP-MS (**Fig. 2 and *SI Appendix*, Fig. S3**). There are significant changes in Fe, Cu, Mn, and Zn with different triggers inducing different changes in metal content compared to cells grown in minimal media. The most notable changes are a decrease in total Cu in ZD, a 2.8-fold increase in Fe and 2.1-fold increase in Mn in GF, and a 4-fold increase in Fe and 2.6-fold increase in Mn in Starv. The increase in total Fe and Mn under serum starvation is surprising given that these metals are largely supplied by the serum, although it could reflect sequestration of these metals under starvation conditions, perhaps from dead cells or upregulation of metal storage proteins. Interestingly, all triggers induce a decrease in Zn compared to cells grown in minimal media.

**Figure 2.**
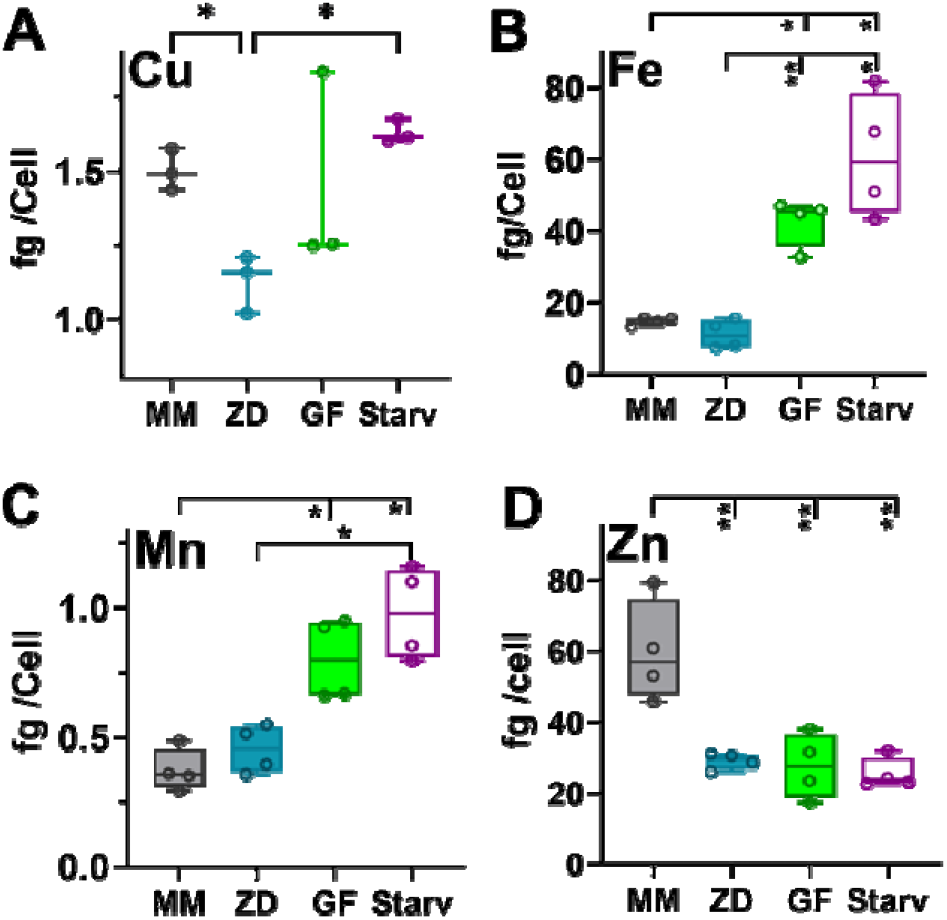
Total metal status differs across cells with different quiescence triggers. ICP-MS analysis of total Cu (A), Fe (B), Mn (C), and Zn (D) from unsorted MCF10A p21-mCitrine geminin-mCherry cells grown in minimal media or treated with different quiescence triggers for 48h and then digested for ICP-MS analysis. All measurements were performed in at least three biological replicates (N ≥ 3). Error bars represent SD. Significance was determined *via* Brown-Forsythe and Welch ANOVA with Dunnett’s T3 multiple comparison test (\**p* < 0.05; ^**^*p* < 0.01; ^***^*p* < 0.001; ^****^*p* < 0.0001).

The changes in total metal content suggest significant changes in metal homeostasis in quiescent cells. Dynamic changes in the labile metal pool can influence cellular processes,(11, 22, 23) so we next characterized the labile metal pool across multiple quiescent states induced by different triggers. To accomplish this, we used small molecule fluorescent probes such as the activity based FIP-1 Fe^2+^ sensor,(24) the binding-based CF4 Cu^+^ sensor,(25) and the M4 Mn^2+^ sensor.(26) We also used a genetically encoded fluorescent sensor for Zn^2+^.(27) We first verified that each probe measured the labile metal pool by treating cells with a chelator or excess metal to demonstrate the probe could detect decreases and increases (respectively) in the metal pool (***SI Appendix*, Fig. S4**). Cells were treated with different quiescence triggers, or grown in MM, for 48 hrs, then stained with the respective small molecule fluorescent probes as described in methods. We observed significant perturbations of the labile metal ion pools upon induction of quiescence, suggesting remodeling of metal homeostasis (**Fig. 3**, and ***SI Appendix*, Table S1**) and different quiescence triggers induced distinct changes in the labile metal ion pool. The most notable differences are a significant increase in labile Cu^+^ in Starv; a significant increase in labile Fe^2+^ in Starv; and a decrease in labile Fe^2+^ in ZD and GF; a significant increase in Mn^2+^ in ZD; and a consistent decrease in Zn^2+^ in all quiescence triggers compared to cells grown in MM. This observation indicates a remodeling of labile transition metal ion pool in quiescence.

**Figure 3.**
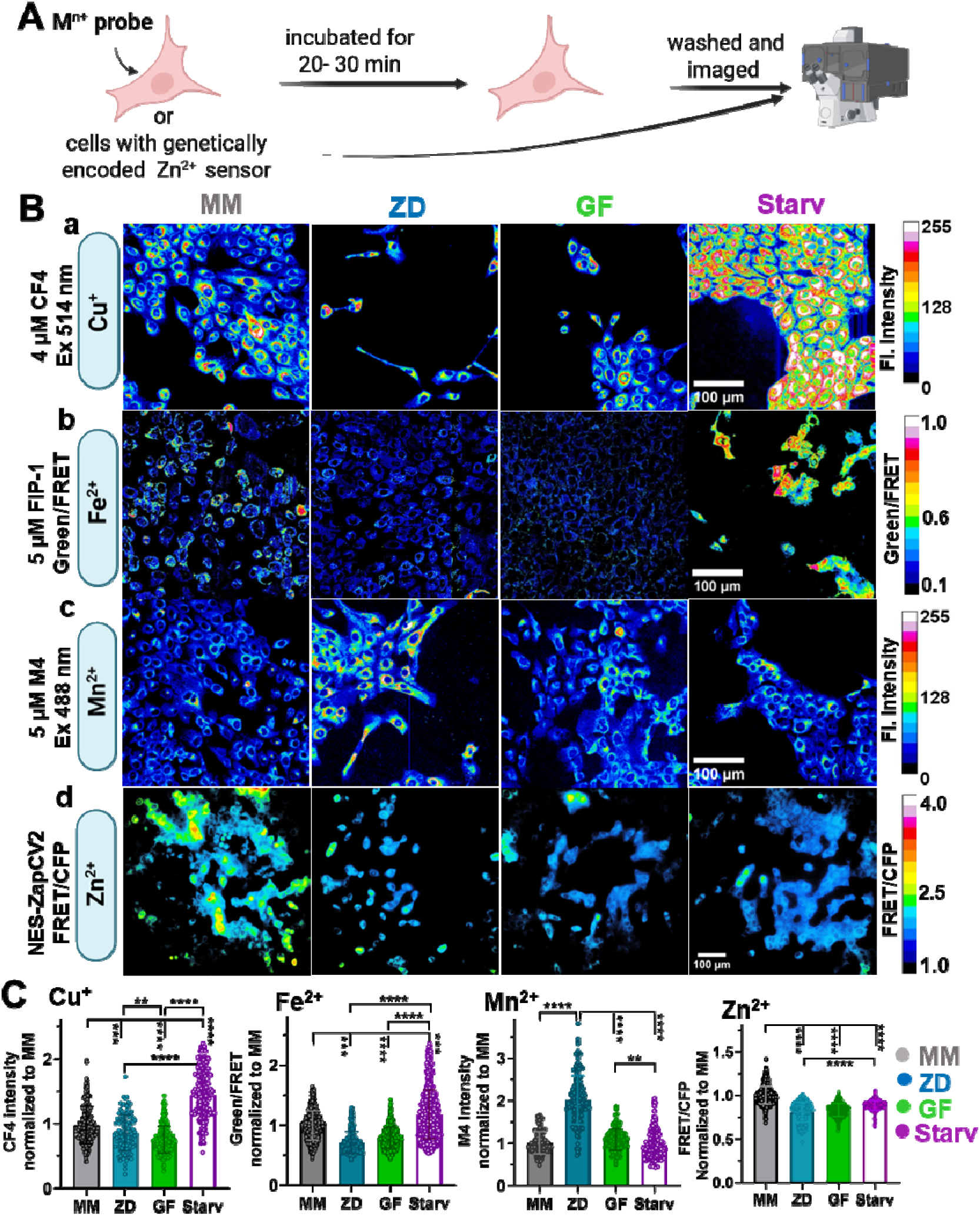
Labile metal ion status in different quiescent cell types. (A) MCF10A cells were treated with specific quiescence triggers for 48h and then incubated with metal ion-specific fluorescent probes in respective media for at least 20 mins. (B) Live cell imaging of cells with (a) turn on based Cu^+^ sensor CF4 (4 µM, 30 min), (b) FRET based Fe^2+^ sensor FIP-1(5 µM, 90 min, FRET ratio: Green/FRET), (c) Turn-on based Mn^2+^ sensor M4 (5 µM, 15 min), and (d) genetically encoded FRET based Zn^2+^ sensor NES-ZapCV2 (FRET ratio: FRET/CFP). Images are pseudo colored and calibration bars indicate signal response ranging from blue (low) to white (high) for each metal probe. Scale bar: 100 µm for (a-d). (C) Normalized (relative to MM) fluorescence intensities or FRET ratios for respective metal ion probes per condition. Each experiment was performed in 96-well plates at least twice with n ≥ 2 wells per experiment. Each dot represents fluorescence response from a single cell and bar plots are color coded to specific quiescence triggers. Statistical analysis was performed and significance was determined *via* Brown-Forsythe and Welch ANOVA with Dunnett’s T3 multiple comparison test (^**^*p* < 0.01; ^***^*p* < 0.001; ^****^*p* < 0.0001). Error bars represent SD.

### There are widespread changes in metal homeostasis and transport genes in quiescence

To investigate whether different quiescence triggers uniquely influence the expression of metal homeostasis and transport genes and gain insight into what genes might be responsible for the changes in total and labile metal ion pools, we identified gene lists for overall metal regulatory genes (2,458 genes) as well as metal binding and homeostasis genes specific to Zn (2,240 genes), and metal transport and homeostasis genes for Cu (57 genes), and Mn (39 genes) (see Methods, ***SI Appendix*, Datasets S2-S3**).

Many metal regulatory genes are differentially expressed in quiescence triggers compared to spontaneously quiescent cells, with Starv and ZD triggers inducing significantly more changes than GF (1354 vs 1221 vs 516 DEGs for Starv, ZD, and GF respectively) (**Fig. 4*A***, and ***SI Appendix*, Figs. S5A-B**). KEGG and Gene Ontology (GO) enrichment analysis of DEGs identified a number of processes that intersect with metal-dependent functions, including metal transport, Fe homeostasis, mitochondrial processes, and oxidative stress (**Fig. 4*B***). 337 DEGs were shared among all three triggers, and these DEGs are enriched in genes related to transport, cellular response to stress, and regulation of cell cycle G2/M phase transition **(*SI Appendix*, Fig. S5C)**, suggesting metal regulatory genes play a critical role in the downregulation of the cell cycle as well as cellular stress in quiescence. Notably, our reanalysis of a previously published dataset for Starv/Spont collected by a different lab showed an 80% overlap with the metal regulatory DEGs (***SI Appendix*, Fig. S5D)**.

**Figure 4.**
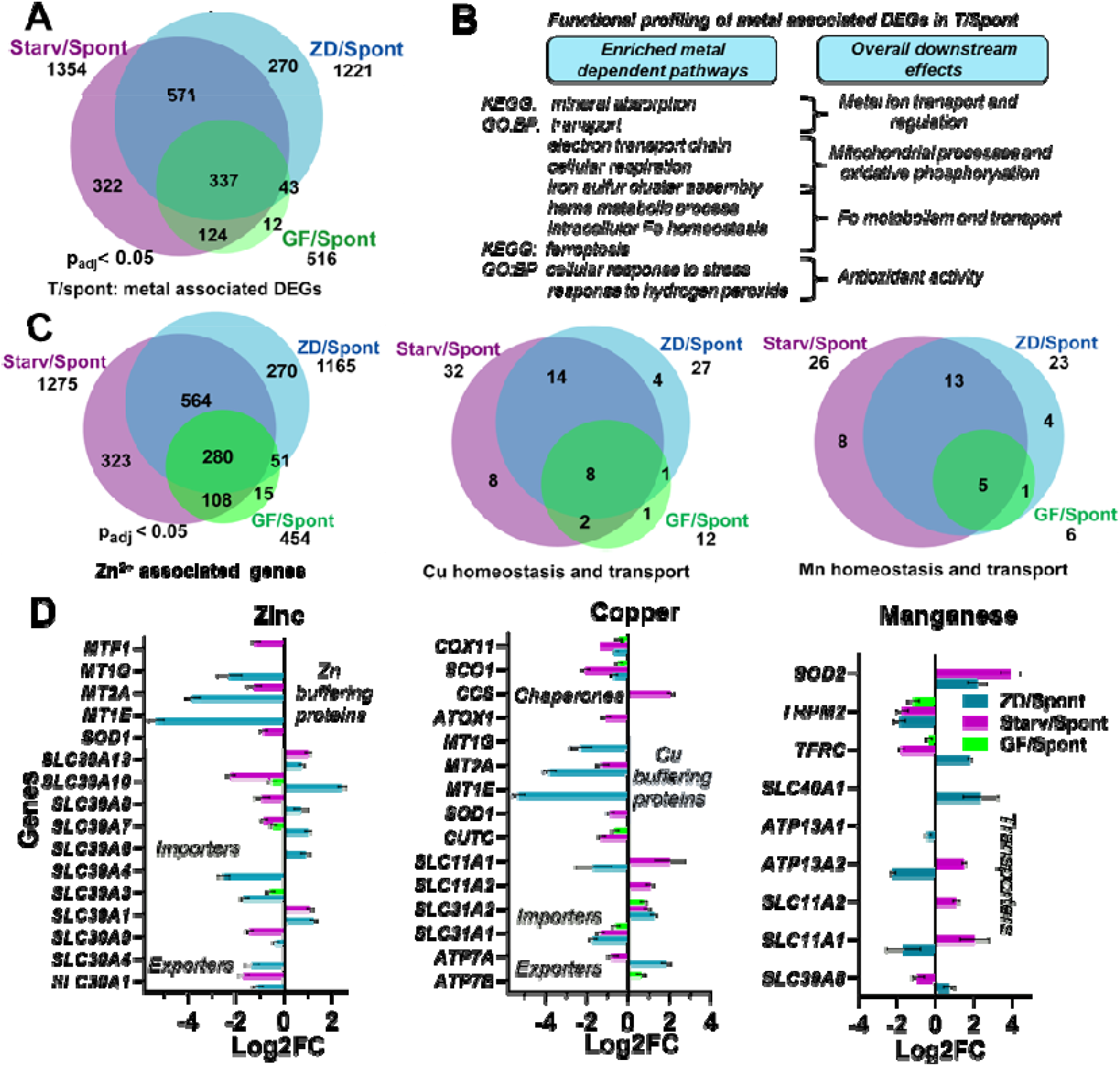
Global remodeling of metal associated genes in quiescence triggers. (A) Venn diagram of differentially expressed metal associated genes from binary comparisons of each quiescence triggers relative to spontaneously quiescent cells (T/Spont, p_adj_ <0.05). (B) Functional analysis of metal associated DEGs in T/Spont revealing significantly enriched metal dependent GO’BP’ and KEGG pathways in quiescence. (C) Venn diagrams of significant zinc, copper, and manganese regulatory DEGs (p_adj_ < 0.05) overlapping with Gene ontology derived gene sets for ‘Zinc ion’ associated, ‘Copper ion homeostasis and transport’, and ‘Manganese ion homeostasis and transport’ in quiescence triggers (See methods for details). (D) Bar plots depict log2foldchange (Log2FC) in gene expression for DEGs (p_adj_ < 0.05) in curated gene sets specific for maintaining critical intracellular labile zinc, copper, and manganese ion levels. Error bars represent standard error in Log2FC by DESeq2. Deep cyan for ZD/Spont, purple for Starv/Spont, and green for GF/Spont.

Quiescent cells exhibit a significant decrease in total and labile Zn, suggesting a universal perturbation of Zn homeostasis. Differential expression of Zn associated genes was widespread across triggers with 1165 genes for ZD/Spont, 1275 genes for Starv/Spont, 454 genes for GF/Spont, and 280 DEGs shared across all triggers (**Fig. 4*C***). Functional analysis revealed common upregulated processes (regulation of RNA biosynthesis) and downregulated processes (transcription, DNA replication, Zn related processes) as well as several processes that were differentially enriched upon different triggers (chromatin organization and protein modification) (***SI Appendix*, Fig. S6)**. In summary, although all quiescent states show decrease in labile and total Zn, each has a distinct signature of altered Zn associated genes.

Changes in intracellular Zn homeostasis can be caused by changes in membrane transport of Zn^2+^ and intracellular storage. Few of these genes are differentially expressed in GF, but in ZD and Starv we observed a strong decrease in expression of genes that encode the Zn storage and buffering proteins metallothioneins (MTs), consistent with the decrease in labile Zn^2+^ in these cells (**Fig. 4*D***). There was also a decrease in many of the Zn export genes *SLC30A1-10* (that encode ZnT transporters), consistent with cellular attempts to limit Zn export to restore Zn levels. There is remodeling of expression of *SLC39A1-14* genes which encode ZIP importers that transport Zn into the cytosol from intracellular organelles and from outside the cell, with some increases and decreases, but *SLC39A1* (ZIP1), the primary Zn importer across the plasma membrane, shows consistent upregulation as an attempt to restore cytosolic Zn under limiting conditions. These results reflect the fact that ZD and Starv quiescent cells display widespread and potentially consequential changes in Zn associated genes, including Zn homeostasis genes that reflect Zn deficiency.

Each quiescence trigger is also associated with distinct changes in Cu homeostasis by remodeling expression of Cu transporters and chaperones which could lead to changes in cellular copper levels **(Fig. 4*C*** and **4*D*)**. In ZD, Cu handling genes showed mixed regulation, with downregulation of genes encoding Cu importers (*SLC31A1* encoding CTR1 and *SLC11A1* encoding NRAMP1) alongside upregulation of the gene encoding the Cu exporter ATP7A, importer CTR2 (*SLC31A2*), and copper chaperones (*COX19, CCS*), consistent with a redistribution of intracellular copper trafficking pathways perhaps to manage the observed decrease in total and labile Cu. Starv displayed a distinct profile marked by reduced expression of Cu-Zn-superoxide dismutase *SOD1* and Cu chaperones (*ATOX1, SCO1*, and *COX17*), and increased expression of membrane importers (*SLC31A2, SLC11A1*, and *SLC11A2*). These changes coupled with downregulation of *ATP7A* exporters suggest a different regulatory pattern that could contribute to the increased total and labile Cu observed in these cells. GF presents another pattern with upregulation of *ATP7B* (encodes a Cu exporter) and downregulation of genes encoding Cu chaperones. GF was associated with a decrease in labile Cu but no change in total Cu.

Manganese transport has been known to use Zn associated ZIP (e.g., *SLC39A14*,(28) *SLC39A* 8)(29) and ZnT transporters (*SLC30A10*),(30) Fe associated *SLC11* transporters and *TFR* receptors,(28, 31) as well as H^+^ and Ca^2+^ channels (e.g., *ATP13A1/2*,(32) *SPCA1*(33)), some of which are differentially expressed upon different quiescence triggers **(Fig. 4*C-D*)**. ZD and Starv both showed upregulation of *SOD2* which encodes a Mn-superoxide dismutase. ZD showed downregulation of *SLC11A1, ATP13A2* and upregulation of *TFRC*. In contrast, *SLC11A1*, and *ATP13A2* are upregulated in Starv/Spont while *TFRC* is downregulated. Moreover, the shared iron exporter *SLC40A1* was up and *ATP13A1* was down only in ZD and the shared zinc importer *SLC39A8* (Zip8) is only downregulated in Starv. Taken together, a trigger specific remodeling of individual metal regulation was noted in quiescence.

### Iron metabolism changes significantly in quiescence

To examine changes in iron metabolism and homeostasis in quiescence, we developed a comprehensive gene list of 139 genes associated with the GO BP terms: heme metabolic processes, iron sulfur cluster assembly, and intracellular Fe homeostasis (***SI Appendix*, Dataset S4)** and examined DEGs in all three triggers (**Fig. 5**, and **Table S2)**. There are widespread changes in genes involved in Fe import, transport, and storage (**Fig. 5*A***), as well as genes associated with intracellular Fe homeostasis, Fe-S cluster biogenesis, and heme metabolism (**Fig. 5*B***)

**Figure 5.**
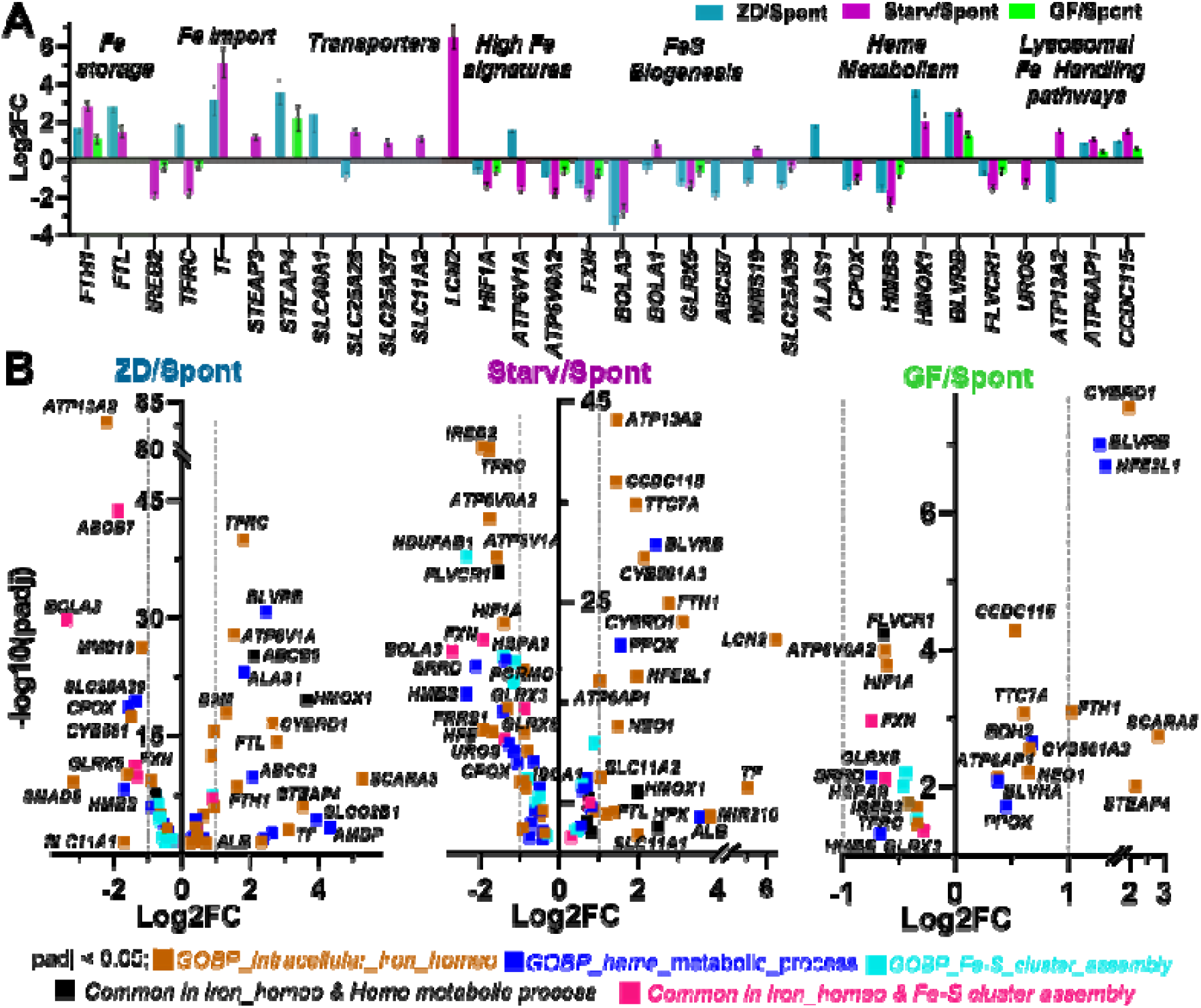
Intracellular iron homeostasis and heme biogenesis in quiescence triggers. (A) Bar plots of Log2FC of DEGs in different quiescence triggers vs spontaneous quiescence based on a curated list of genes associated with intracellular iron homeostasis, FeS cluster biogenesis, heme biosynthesis enzymes, and heme metabolism genes (Table S2). Error bars represent standard error in Log2FC by DESeq2. (B) Volcano plots of DE genes (padj < 0.05) associated with GO BP ‘intracellular iron homeostasis’ (GO:0006879), GO:0016226 ‘Iron Sulphur clusters assembly’, GO:0042168 ‘Heme metabolic Process’ in each binary T/Spont comparison. Each dot represents a DE gene. DEGs within dotted line have Log2FC <|1|.

ZD shows upregulation of iron uptake and ferrireductases that facilitate iron import (*TFRC, CYBRD1, STEAP4, SCARA5*, and *SLCO2B1*), while Starv shows signs of decreased uptake (downregulation of *TFRC* and *IREB2*). The gene encoding iron responsive element binding protein 2 (*IREB2*) has been previously implicated in the regulation of *TFRC* and ferritin (*FTH1, FTL*) expression and *ireb2* knockdown in mice showed increased ferritin with concomitant decrease in *TFRC* expression in brain.(34) Consistent with this regulatory role, we observed downregulation of *IREB2* in Starv accompanied with expected modulation of target genes. Starv also shows signs of Fe remodeling due to high intracellular Fe (upregulation of *LCN2*, and downregulation of *HIF1A, ATP6V1A*, and *ATP6V01B*) and altered lysosome and Fe handling due to upregulation of *ATP13A2, ATP6AP1*, and *CCDC115*, perhaps due to increased lysosomal degradation or mitophagy. ZD, in contrast, exhibits significant downregulation of *ATP13A2* and upregulation of iron uptake.

Digging further into Fe transport, we looked at canonical Fe transporters including a plasma membrane Fe importer (*SLC11A2* which encodes DMT1), mitochondrial Fe importers (*MFRN1* and *MFRN2: SLC25A37/28*), and the sole Fe exporter (*SLC40A1* which encodes the ferroportin protein) (**Fig. 5*A***). Surprisingly, *SLC11A2, MFRN1*, and *MFRN2* were all increased in Starv, despite these cells having high labile Fe. On the other hand, *SLC40A1* was increased in ZD despite these cells having low labile Fe, and *MFRN2* was decreased which would reduce Fe import into mitochondria perhaps contributing to compromised Fe-S cluster biosynthesis in mitochondria.

ZD and Starv are characterized by downregulation of Fe-S cluster biogenesis (*FXN, BOLA3, GLRX5*), but ZD shows additional downregulation of *MMS19, SLC25A39*, and *ABCB7*, suggesting more widespread suppression of Fe-S cluster biogenesis (**Fig. 5*B***). Both ZD and Starv also show dysregulation of heme metabolism with changes in biosynthesis (*ALAS1, HMBS, UROS, CPOX, PPOX*), transport (*ABCB6, FLVCR1*), and degradation (*HMOX1*) (**Fig. 5*B***). This is perhaps not surprising given that 48% and 17% of the Fe binding proteins in human genome are heme binding and Fe-S cluster proteins, respectively (35) and mitochondrial Fe-S cluster defects can disrupt heme metabolism.(36) Finally, both ZD and Starv show an increase in iron storage genes (ferritin genes *FTH1*, and *FTL*).

Fe regulatory genes appeared less perturbed in GF, despite exhibiting elevated labile and total Fe. We observed genes related to Fe import to variably expressed (*IREB2, TFRC, CYBRD1* and *STEAP4*), while the ferritin receptor *SCARA5*, ferritin subunit *FTH1*, and *NFE2L1* are upregulated. *NFE2L1* is a master transcriptional regulator of the labile Fe pool which increases ferritin expression to promote Fe storage, suggesting an increase in Fe storage capacity with this trigger.

Taken together these results indicate significant perturbation of Fe regulatory, homeostasis, and transport genes with different signatures for different quiescence triggers. These findings are consistent with a broader role of Fe homeostasis in regulating proliferation across stem and immune cells, where quiescent hematopoietic stem cells tightly restrict the labile Fe pool to prevent age-related functional decline associated with even mild Fe loading, (37) and where Fe availability similarly controls native CD4 T cell quiescence, with excess Fe impairing mitochondrial integrity and glycolytic function.(38)

### Altered metal homeostasis is associated with changes in oxidative phosphorylation and antioxidant activity

Functional profiling of the 2458 metal homeostasis, transport, and dependent DEGs in quiescence triggers revealed enrichment of mitochondrial processes and oxidative stress (**Fig. 4*B***). To gain further insight into how these biological processes might be altered in the different quiescent states, we selected BP oxidative phosphorylation (OXPHOS, 148 genes) and BP antioxidant activity (58 genes) for further analysis (see Methods). ZD and Starv show large changes in OXPHOS genes (43% and 56% of the 148 genes, respectively) compared to GF (13%) (**Fig 6*A***). A large fraction of these genes are metal associated (47 out of 63 DEGs for ZD, 60 out of 83 DEGs for Starv, 11 out of 19 DEGs for GF). Upregulated genes that are common for all three triggers include *PINK1*, responsible for mitophagy induction, *NUPR1*, a stress-induced transcription factor that promotes quiescence, and *ABCD1*, a fatty acid transport protein involved in lipid metabolism. Common downregulated genes include cell cycle regulators *CDK1* and *CCNB1*.

**Figure 6.**
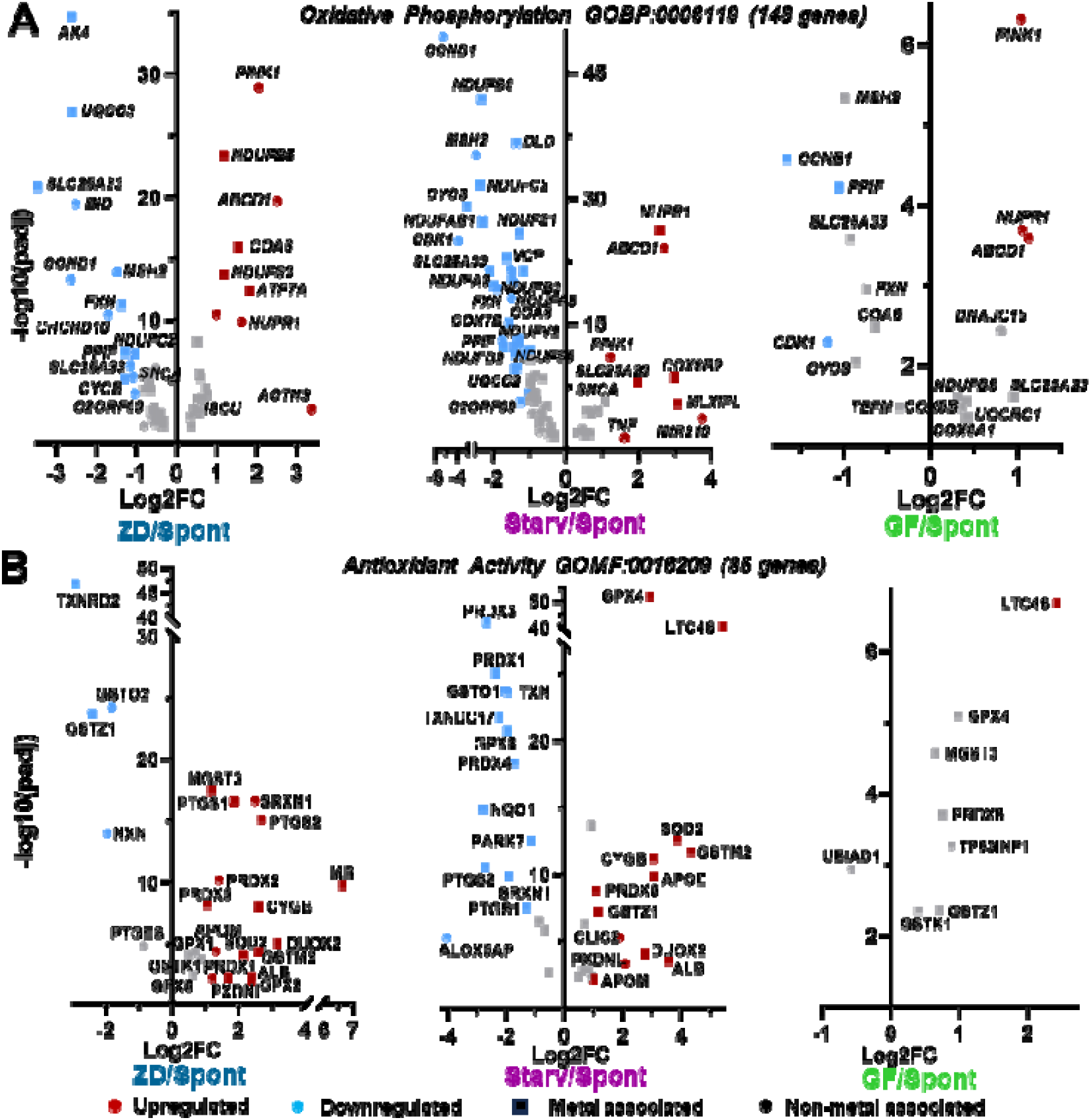
Altered mitochondrial oxidative phosphorylation and cellular antioxidant activity associated gene landscape in quiescence triggers. (A) Volcano plots of the DE genes (p_adj_ <0.05) associated with GO:0006119 (BP_Oxidative_Phosphorylation) in ZD/Spont, Starv/Spont, and GF/Spont. Squares represent metal-associated genes. (B) Volcano plots of the DE genes (p_adj_ <0.01) associated with GO:0016209 (MF_Antioxidant_Activity) in ZD/Spont, Starv/Spont, and GF/Spont. Each dot represents a DE gene and grey dots denote DEGs with Log2FC <|1| for (A) and (B).

ZD and Starv show significant changes in genes related to OXPHOS compared to GF, but specific changes in OXPHOS genes indicate that ZD and Starv perturb the process differently (**Fig. 6*A***). ZD is characterized by upregulation of *NDUFS3* and *NDUFB5* which would promote Complex I stabilization, as well as *COA6* which is involved in Cytochrome C maturation and suggests Complex IV assembly, and finally *ATP7A* which is involved in Cu transport and necessary for Complex IV. Downregulated genes include some assembly factors (*UQCC3*), metabolite transporters (*SLC25* family), and a maturation factor (*CYC3*). These changes suggest adaptive remodeling of mitochondria, with selective stabilization of some functions and trimming of others. Conversely, Starv is characterized by broad suppression of Complex I (downregulation of 7 *NDUF* catalytic and accessory subunits), along with a decrease in other respiratory chain components (*CYCS, COX7B, COA6*) which would reduce complex IV maturation and decrease electron transport capacity. Finally, Starv shows a decrease in *FXN* which would decrease Fe-S cluster assembly, and a decrease in *DLD*, an E3 component of multiple dehydrogenase complexes. A decrease in *DLD* could lower flux through pyruvate dehydrogenase, lower α-ketoglutarate dehydrogenase activity, and lower NADH production. The upregulation of *COX6A1/COX6B2* suggest some selective balancing of cellular respiratory subunits. The changes in Starv suggest broad decrease in mitochondrial function consistent with decreased energy capacity.

There is also a striking difference in the pattern of DEGs involved in antioxidant activity that suggests zinc deficient quiescent cells upregulate antioxidant activity whereas cells driven into quiescence by serum starvation show up and down regulation of antioxidant genes (**Fig. 6*B***). GF showed only 8 DEGs, with only *LTC4S* showing a log2FC > 1, suggesting very little perturbation of antioxidant activity in GF. In contrast, ZD has 25 DEGs out of 85 genes and Starv has 35 DEGs out of 85 genes. We found that 20 out of 25 DEGs for ZD/Spont and 32 out of 35 DEGs for Starv/Spont are metal associated. ZD is characterized by upregulation of systems that detoxify reactive oxygen species (ROS), including *sod2*, a Mn-dependent superoxide dismutase that detoxifies superoxide; peroxiredoxins, *PRDX1/2/5* that detoxify hydrogen peroxide; *SRXN1*, involved in recycling of hyperoxidized peroxiredoxins; *GPX2* involved in glutathione dependent peroxide removal; and *GSTM1*, a glutathione transferase involved in detoxification of electrophilic compounds. Also, upregulated are *PTGS1/2* which convert arachidonic acid to prostaglandins and act as central mediators of inflammation that are activated by oxidative stress (**Fig. 6*B***). While there are few downregulated genes, the most striking is *txnrd2* a selenium-dependent thioredoxin. It is possible that zinc deficiency alters selenium; this metal was not evaluated by ICP-MS.

In contrast to broad upregulation of antioxidant systems, Starv shows more balanced remodeling (**Fig. 6*B***). This includes upregulation of the Mn-dependent superoxide dismutase (*SOD2*), genes involved in glutathione conjugation (*GSTM2, GSTZ1*), and downregulation of a number of peroxiredoxins (*PRDX1/3/4*), as well as *SRXN1* and *PTGS1/2* which were upregulated in ZD. Interestingly, the two most significantly upregulated genes are *GPX4* which uses glutathione to protect against lipid peroxidation and ferroptosis but not generic ROS production, and *LTC4S*, an enzyme which consumes glutathione for the biosynthesis of leukotrienes. In summary, ZD appears to be characterized by broad active suppression of oxidative stress, while Starv suggests a more mixed pattern.

## Discussion

Metal ions are essential to life. Bioinformatics estimates suggest that up to 40% of enzymes use a metal as a cofactor for catalysis.(39) Analysis of sequence and structural homology led to the prediction that up to 10% of human proteins (∼ 2500) contain zinc binding motifs,(40) with hundreds of additional proteins predicted to bind iron, copper, calcium, manganese, and magnesium.(35, 41) Modern proteomics approaches that identify metalloproteins experimentally often suggest the number of metalloproteins is even higher.(42–44) Furthermore, evidence is emerging that even when proteins don’t contain a predicted metal binding site, function can be modulated by metal binding.(13, 44) While metal ions are clearly necessary to populate thousands of metalloproteins and enzymes, how cells cope with changes in metal availability, and more specifically how metal availability affects proliferation and quiescence is less well defined. Yet serum starvation is one of the most common triggers of quiescence and serum starvation involves depriving cells of essential metals, as well as other nutrients and growth factors. Multiple studies have shown that Fe regulates quiescence and cell fate of hematopoetic stem cells,(45) CD4 T-cells,(46) and Fe deficiency impairs muscle (47) and mesenchymal (48) stem cell proliferation. We have shown that there is a pulse of Zn in early G1 in the mammalian cell cycle,(49) that Zn deficiency induces quiescence,(5) and that a short (2 hr) pulse of low Zn stalls mammalian cells in S-phase and induces transient cell cycle withdrawal in daughter cells,(50) indicating Zn plays important regulatory roles in the mammalian cell cycle. While there is clear evidence that metal availability can play a regulatory role in the proliferation-quiescence axis, how metal homeostasis is altered in quiescence has not been systematically explored. The purpose of this study was to examine the transcriptional changes in zinc deficiency-induced quiescence and compare to transcriptome remodeling induced by other triggers to identify core transcriptional changes common to all quiescent states and those that are unique to zinc deficiency. Further, we sought to evaluate whether transition metal homeostasis is universally remodeled in quiescence, and if so how. Finally, we sought to provide insight into the metal homeostasis genes that are altered in quiescence and what cellular pathways and processes could be affected as a result.

Despite being a limited nutrient perturbation compared to serum starvation, zinc deficiency induces widescale changes in gene expression. Some of the gene expression changes are consistent across all triggers, indicating ZD-induced quiescence is similar to other triggers in downregulating cell cycle processes, mTOR signaling, and DNA replication and upregulating autophagy and lysosomal processes (**Figure 1*F***). But the transcriptional program of ZD quiescence is distinct with 29% (2,584 genes) of the total DEGs unique to ZD. Even when there are shared altered pathways between quiescent states, ZD presents a distinct signature. Here, we report the first comprehensive picture of transcriptional remodeling in ZD induced quiescent cells. The ZD quiescent state is characterized by a remodeling of mitochondrial oxidative phosphorylation and upregulation of cellular antioxidant activity, chromatin organization, and protein ubiquitination processes. The upregulation of antioxidant genes in ZD quiescence (**Fig. 6*B***) is consistent with previous observations on increased peroxiredoxin and glutathione transferase levels in zinc deficient *C. reinhardtii*,(51) upregulated *TSA1* peroxiredoxin in zinc deficient yeast,(52) and increased SOD1/2 activity in zinc deficient 3T3 cells,(53) and aligns with well-established antioxidant role of Zn in maintaining redox homeostasis.(54, 55)

Quiescent cells are known to conserve energy by repressing mitochondrial OXPHOS,(56–58) upregulating antioxidant defenses,(5, 11, 50) and reducing translation (59, 60) via intron retention (61), as documented across quiescent epithelial,(2, 58) fibroblast,(4) immune,(38) and stem cells.(45, 56) Consistent with this, serum starved quiescent cells globally repress mitochondrial *OXPHOS* gene expression and adapt to stress via a balanced antioxidant transcriptional response (**Fig. 6**). In contrast, ZD quiescent cells remodel the OXPHOS pathway in a structured way (**Fig. 6*A***) suggesting a distinct regulatory mechanism. Interestingly, the upregulation of chromatin organization in ZD is consistent with previous observations that zinc alters chromatin accessibility in MCF10A cells (62) and with the known role of epigenetic remodeling including histone methylation (63) and alteration in chromatin organization (6, 64) in preserving quiescence reversibility. Finally, Zn’s structural role in E3 ligase function (65) makes the upregulation of protein ubiquitination in ZD quiescence consistent with the E3-ligase mediated proteasomal degradation for nutrient recycling in quiescent cells.(16, 66–68)

All quiescent states were associated with significant remodeling of metal homeostasis and concomitant changes in the total and labile metal pools. Quiescent cells are characterized by a decrease in both total and labile Zn, regardless of the trigger, compared to cells grown in minimal media, suggesting that decreased Zn may be a signature of quiescence. Individual quiescence triggers also showed significant perturbation of Zn-associated genes, with 1165 DEGs for ZD/Spont, 1275 DEGs for Starv/Spont, and 454 DEGs for GF/Spont out of 2240 Zn associated genes in the list. A striking proportion are inversely regulated between conditions (***SI Appendi*x, Fig. S7)**. 37% of shared Zn associated DEGs between ZD and Starv are reciprocally regulated (***SI Appendi*x, Fig. S7B**), as are 17% of shared Zn associated DEGs between GF and ZD). This reciprocal regulation suggests distinct regulation of Zn associated genes in quiescence states despite similar changes to total and labile Zn.

We observed widespread but distinct changes in Fe, Cu, and Mn, and associated metal transport, homeostasis, and dependent genes in different triggers. Analysis of metal transport and homeostasis genes revealed genes that could be related to altered metal pools, but we could not distinguish whether changes in gene expression are a cause or consequence. The Irving-William series, which describes the relative stability of transition metal complexes (Mg^2+^ < Mn^2+^ < Fe^2+^ < Co^2+^ < Ni^2+^ < Cu^2+^ > Zn^2+^), is frequently invoked to explain intracellular metal ion availability, where more competitive metals (Cu, Zn) are maintained at far lower concentrations than Mn and Fe to prevent mis-metallation of proteins.(69) However, compartment-specific metal partitioning can further enforce metal selectivity as shown by differential metalation in CucA and MncA cupins in cyanobacterium.(70) Crosstalk among metal pools is therefore expected given the promiscuous nature of shared ZIPs, ZnTs, DMT1 and TfR-1 transporters for Zn^2+^, Fe^2+^ and Mn^2+^ as observed in metal dependent growth perturbations across Ca, Cu, Fe, Mn, and Zn in *S. cerevisiae* (71) and Zn/Fe/Mn interdependence in mammalian systems.(29, 66–68) For example, knockout of the Mn exporter SPAC1 or external Mn treatment in HAP cells reduces MT, ZnT1 and overall Zn levels which get rescued upon ZnT10 expression.(33) Further, mutation of the ZIP8 transporter causes severe Mn depletion.(72) This close crosstalk between Zn and Mn homeostasis might underlie the low Zn but high labile Mn pool in ZD quiescence. Knockout of the Cu exporter *ATP7A/7B* increased Cu but did not change Zn levels in HAP cells.(33) Mn overload in both *slc39a14*^-/-^ mice (73) and *slc39a14*^U801^ mutant zebrafish (28) showed reciprocal metal dysregulation, where elevated Mn decreases Zn uptake while also enhancing Fe accumulation through upregulation of Fe-transporters DMT1 and TfR-1 (73) and TfR1B expression (28). Furthermore, ZIP13 overexpression reduces cytosolic Fe but knockout increases total Fe while depleting mitochondrial and lysosomal labile Fe in MEF cells.(74) These studies highlight the interconnected nature of the cellular metal homeostasis.

The changes in total and labile metal pools and corresponding transcriptional changes in Fe, Zn, Cu and Mn regulatory genes suggest perturbation in one specific metal pool during quiescence has broader downstream consequences on the cellular metal landscape. This may cause the distinct regulation of major redox metal dependent processes including oxidative phosphorylation and heme metabolism across quiescence states. Our study provides a first report on metal homeostasis as a massively remodeled axis in response to quiescence reiterating the importance of being attentive to the sensitivity of metal homeostasis to nutrient availability when studying cellular quiescence. Our findings on changes in transition metal pools and metal transcriptomics serve as an important reminder that metal homeostasis remodeling is likely a fundamental and underappreciated component of cellular quiescence.

## Materials and Methods

### Cell culture

MCF10A wildtype cells were procured from ATCC. MCF10A H2B-HaloTag, and MCF10A NES-ZapCV2-H2B-HaloTag were generated by the Palmer laboratory following previous protocol.(5) The MCF10A p21-mCitrine geminin mCherry cell line was generated by the Spencer laboratory at CU Boulder(2). All MCF10A cell lines were maintained in full growth medium (FGM) consisting of DMEM/F12 medium supplemented with 5% horse serum, 1% pen/strep antibiotics, 20 ng/mL epidermal growth factor (EGF), 0.5 μg/mL hydrocortisone, 100 ng/mL cholera toxin, and 10 μg/mL insulin. Cells were passaged with 0.05% trypsin-EDTA. To remove excess Zn^2+^ from horse serum and insulin and to generate the defined minimal medium (MM), we incubated serum and insulin solutions with Chelex-100 for 12 h, followed by sterile filtration to remove Chelex-100 resin. Minimal media (MM) consisted of 50:50 Ham’s F12 phenol red free/FluoroBrite DMEM with 1.5% Chelex 100-treated horse serum, 10 μg/mL Chelex 100-treated insulin, 1% pen/strep antibiotics, 20 ng/mL EGF, 0.5 μg/mL hydrocortisone, and 100 ng/mL cholera toxin. ZD media was generated by adding 3 μM TPA to the minimal media. GF media was comprised of MM without EGF and insulin. Serum starvation media was comprised of MM without serum, EGF, and insulin. Cells were grown in a humidified incubator at 37°C and 5% CO_2_. Cell lines were routinely tested and confirmed to be mycoplasma negative by PCR.

The detailed procedures for cell culture, imaging, and analyzing RNA sequencing data from quiescent cells along with full experimental details including protocols, and data analysis parameters, are provided in *SI Appendix, Materials and Methods*.

## Data Availability

Datasets will be deposited in Gene Expression Omnibus (GEO) database and available in the *SI Appendix*.

## Acknowledgments

We thank the University of Colorado Boulder Cell Culture Core Facility (RRID:SCR_018988) and Flow Cytometry Core facility, specifically Theresa Nahreini for assistance with cell culturing and sorting. We would thank the BioFrontiers Advanced Light Microscopy Core (RRID: SCR_018302) for imaging resources and BioFrontiers computing Core for providing High Performance Computing resources. ICP-MS measurements were performed in the OHSU Elemental Analysis Core (RRID: SCR_022746) with partial support from NIH (S10RR025512). We thank Prof. Chris Chang for helpful discussions and support on metal imaging experiments. We thank Prof. Sabrina L. Spencer at University of Colorado Boulder for providing the MCF10A p21-mCitrine geminin-mCherry cell line. We thank BioFrontiers Institute Next-Gen Sequencing Core Facility, Genomics Shared Resource at University of Colorado Anschutz Medical Campus (RRID: SCR_021984) and the Cancer Center Support Grant (P30CA046934) for performing RNA-sequencing on our samples. A. D. received support from Department of Atomic Energy, Government of India (Project Identification No. RTI4015) for development of M4 probe at TIFR Mumbai. We would like to acknowledge NIGMS MIRA R35 GM139644 (A.E.P.) for generous financial support.

