## Supplemental appendix for "Metal homeostasis is remodeled in response to different quiescence triggers"

**This PDF file includes:**

Supporting text

Figures S1 to S6

Tables S1 to S2

Legends for Datasets S1 to S4

SI References

**Other supporting materials for this manuscript include the following:**

Datasets S1 to S4

Supporting Information Text

**Materials and Methods**

**Fluorescence-activated cell sorting (FACS) to isolate quiescent cells**

To identify a treatment time window with a maximum percentage of quiescent cells, we treated MCF10A p21-mCitrine geminin-mCherry cells with MM, ZD, GF, and starvation media for 24 h, 48 h, and 72 h. After treatment, cells were trypsinized and pelleted. Cell pellets were carefully broken down by pipetting and the cell solution in 1X phosphate buffered saline (PBS) prepared in chelexed MilliQ water was then filtered through a 0.45 µm filter to ensure a dispersed cell suspension in devoid of clumps. Cells were then immediately run on a BD Celesta and sorted based on the mCitrine channel (ex 488nm, em 530/30 nm) and mCherry channel (ex 561nm, em 610/20 nm) signal intensities. MCF10A wild-type cells served as a negative control. After setting up gates for single cells, the quiescent population of cells was identified based on high p21-mCitrine signal intensity and low geminin-mCherry signal intensity as described in previous literature,(1) followed by setting up a collection gate for quiescent cells as shown in Supporting **Fig. S1*A***. Sorted quiescent cells were collected in respective quiescence triggering media and processed for RNA-sequencing.

**Elemental analysis of total metals via ICP-MS**

To measure the total Mn, Fe, Co, Cu, Ca, P, S, and Zn, ICP-MS was performed on MCF10A wild-type cells grown in MM and MCF10A p21-mCitrine geminin-mCherry cells grown in ZD, GF, and starvation media for 48 hours. After 48 hours, cells were spun down and washed twice with 1X PBS prepared in Chelex-100 treated water. Cells were then counted four times before digestion and an average value was used for normalization with cell counts. Cell pellets containing 0.9-15 million cells were vortexed for 3 mins in 100 µL of elemental grade TraceSELECT Ultra nitric acid and digested for 2 h at 90 ^o^C on a heat-bath with occasional swirling of the tubes. After digestion, the mixture was diluted with 900 µL of 1% HNO_3_ in chelexed MilliQ water. All ICP-MS measurements were performed in the OHSU Elemental Analysis Core with partial support from NIH (S10RR025512) and results were normalized to the average cell counts. ICP-MS measurements were performed in N ≥3 biological replicates. Outliers are excluded from analysis. Plots and figures were generated using GraphPad Prism and Adobe Illustrator CS, respectively.

**Live cell imaging**

MCF10A wildtype or MCF10A NES-ZapCV2 H2B-HaloTag or MCF10A H2B-HaloTag cells were plated at a density of 4000-7000 cells/well in FGM in glass-bottom 96-well plates (P96-1.5H-N, Cellvis, Mountain View, CA) and the media was changed to ZD, MM, GF, and starvation conditions 24 hr after plating. Cells were left in those conditions for 48 hours, and they were treated with respective metal-selective fluorescence probes, washed and replenished with MM or respective quiescence media conditions before imaging. Images were acquired on a Nikon Spinning Disk confocal microscope with Andor iXon 888 ultra (EMCCD camera) or Nikon Ti-E High Content Analysis inverted microscope with a Lumencor SPECTRA X light engine (Lumencor, Beaverton, OR) and Hamamatsu Orca FLASH-4.0 V2 cMOS camera (Hamamatsu, Japan) at BioFrontiers Institute’s Advanced Microscopy Core (RRID:SCR_018302). Excitation and emission parameters are available in ***SI Appendix*, Table S1**. Images were processed in ImageJ and pseudo color was used for representation.

***Labile Cu^+^ imaging with turn-on based CF4 sensor***

Stock solutions of 5 mM CuCl_2_ and 50 mM bathocuproinedisulfonic acid (BCS) were freshly prepared in chelexed MilliQ water. The final concentration in each well was maintained at 100 µM CuCl_2_ and 500 µM BCS in minimal media. Cells were incubated with 100 µM CuCl_2_ or 500 µM BCS in minimal media for 1 h in a 37 ^o^C incubator. After 1h, cells were washed twice with 1X PBS. Primary stocks for *turn-on* based Cu^+^ sensor, CF4 (1 mM) were made in cell culture grade DMSO and were kept in -20 ^o^C for long-term storage. Cells in all wells, including different quiescence conditions, were incubated with 4 µM CF4 in respective media for 20 minutes at 37 ^o^C. Cells were again washed twice with 1X PBS before imaging. Live cell imaging was performed with a Nikon spinning disc confocal microscope using a 20X objective at 37 ^o^C with 80% humidity. After imaging, the CF4 fluorescence intensity was measured from cells by selecting a region of interest from the confocal images of cells and no background correction was performed.

***Labile Fe^2+^ imaging with FRET based FIP-1 sensor***

Primary stock solutions of 10 mM ferrous (II) ammonium sulphate (FAS) and 10 mM desferrioxamine (DFO), an iron chelator, were prepared freshly in degassed (30 min N_2_ bubbled) chelexed MilliQ water. Primary stock of 1 mM FIP-1 was prepared in cell culture grade DMSO and was stored at -20 °C. MCF10A H2B-HaloTag cells incubated with 1 mM FAS for 90 min were washed, followed by treatment with 5 µM FIP-1 for 90 min, washed twice with 1X PBS, and then imaged. Green/FRET ratio reflects the labile Fe^2+^ concentration inside cells. To measure a decrease in the labile Fe^2+^ pool, cells were pretreated with 250 μM DFO for 4h, then stained with FIP-1 for 90 min. Cells in different quiescence conditions were treated with FIP-1 for 90 mins and 10 nM JF669 for 15 mins at 37 °C and washed twice with 1X PBS before imaging. Live cell imaging was performed with a Nikon spinning disc confocal microscope using a 20X objective at 37 ^o^C with 80% humidity. Images were nuclear segmented, and the Green/FRET ratio was calculated by dividing the background corrected green channel with the background corrected FRET channel.

***Labile Mn^2+^ imaging with turn-on based M4 sensor***

A stock solution of 5 mM MnCl_2_^.^6H_2_O was freshly prepared in chelexed MilliQ water. MCF10A wildtype cells were incubated with 25 µM MnCl_2_ in water for 1h and 50 µM TPEN (25 mM primary stock in DMSO) in 1X modified Thomson’s buffer (HEPES (20 mM), NaCl (146 mM), KCl (5.4 mM), MgSO_4_ (0.8 mM), KH_2_PO_4_ (0.4 mM), Na_2_HPO_4_ (0.3 mM) and glucose (5.5 mM), pH 7.4) for 40 min at 37 °C. Cells were washed thrice with 1X buffer before imaging. A primary stock of M4 was made in water and kept at 4 ^o^C for short-term storage. Cells in all wells, including different quiescence conditions, were incubated with 5 µM M4 in 1X Thompson buffer for 15 minutes at 37 ^o^C. Cells were again washed three times with buffer before imaging. Live cell imaging was performed with a Nikon spinning disc confocal microscope using a 20X objective at 37 ^o^C with 80% humidity. After imaging, the M4 fluorescence intensity was measured from single cells by selecting a region of interests from the cytoplasm of each cell and no background correction was performed.

***Labile Zn^2+^ imaging with genetically encoded cytosolic ZapCV2 sensor***

Quiescent MCF10A NES-ZapCV2-H2B-HaloTag cells in ZD, MM, GF, and starvation conditions (48h) were treated with 10 nM JF669 HaloTag ligand dye for 15 min at 37 °C, washed and imaged on a Nikon Ti-E High Content analysis inverted microscope using either 10X or 20X objective. FRET ratio images were generated in ImageJ by independently background correcting the FRET (CFP_ex_YFP_em_) and CFP (CFP_ex_CFP_em_) channels following thresholding, then dividing the background corrected FRET channel by the background corrected CFP channel using the ImageJ ‘image calculator’ function. FRET ratio from cells expressing NES-ZapCV2 was measured by selecting region of interest from the ratio images of single cells. Cells not expressing NES-ZapCV2 were excluded from calculations.

**Image analysis and statistical tests**

Each experiment was performed in 96-well plates at least two times with n ≥ 2 wells per experiment. Image analysis for Fe^2+^ was performed on ImageJ with nuclear segmentation using H2B-HaloTag nuclear dye JF669-HaloTag ligand (5 nM or 10 nM) and for Zn^2+^ sensors fret ratio was measured by selecting ROIs from background corrected ratio images of single cells expressing NES-ZapCV2. For images with CF4 and M4 probes, fluorescence intensity was measured by selecting ROIs from each cell using ImageJ and no background correction was performed. Plots were made in GraphPad Prism 11. Plots show the normalized fluorescence intensity or FRET ratios of respective probes for each condition compared to the average values of the MM condition. Statistical analysis was performed, and significance was determined *via* Brown-Forsythe and Welch ANOVA with Dunnett’s T3 multiple comparison test (**p* < 0.05; ^**^*p* < 0.01; ^***^*p* < 0.001; ^****^*p* < 0.0001). Error bars represent SD.

**Sample preparation and RNA-sequencing**

MCF10A p21-mCitrine geminin-mCherry cells grown in ZD, GF, and MM media for 48h were FACS sorted for high p21-mCitrine signal and low geminin-mCherry signal. 500,000 sorted quiescent cells were pelleted, washed with 1X PBS twice, and flash-frozen with liquid N_2_. RNA extraction was performed on all samples at once after one freeze-thaw cycle via Qiagen RNEasy mini kit, and DNase I treatment was done on the spin-column. RNA quality was checked via Tape Station, and library prep was done with an Illumina TruSeq LT kit, which included polyA selection. Single-end 75-base with 40M base pair read/sample sequencing was done on a NextSeq 2.1.0 Illumina machine. The tapeStation run, library prep, and sequencing were performed by the BioFrontiers sequencing core. In a separate experiment, cycling unsorted MCF10A wildtype cells and around 1 million sorted MCF10A p21-mCitrine geminin-mCherry cells with high p21-mCitrine signal and low geminin-mCherry signal after treatment with serum starvation media for 48 h were collected. RNA was extracted from these samples via Qiagen RNEasy mini kit, and DNase I treatment was done on the spin-column. and the quality was checked through via RNA-TapeStation. In this experiment, the tape station run, library prep, and paired end 150 bases with 40M base pair read/sample sequencing was done on a NovaSeq X Plus machine by the Genomics Shared Resource at University of Colorado Anschutz facility. All the experiments were done in four replicates.

**RNA-Seq data analysis pipeline**

Analysis of raw RNA sequencing data was performed using BioFrontiers IT HPC computing services. Analysis for single end (Spont, ZD, and GF) and paired end (Cycling and Starvation) samples were performed separately. For single end reads, we performed data quality assessment with FastQC (v.0.11.5) (2) and used Trimmomatic (v.0.36) (3) to perform TruSeq2-SE adapter trimming and trim off the first 10 bases while discarding reads under 36 bases*.* Reads were then mapped and processed with HISAT2 (v.2.1.0) and Samtools (v.1.8) (4, 5) using default quality settings. Read counting was performed over hg38gtf indexing file from NCBI with featureCounts (R v.4.3.1, Rsubread v.1.6.2 (6, 7)) using the following parameters: *GTF.featureType="exon", useMetaFeatures=TRUE, allowMultiOverlap=FALSE, largestOverlap=FALSE, countMultiMappingReads=FALSE, strandSpecific=2*. For paired end reads, after FastQC analysis, we used Trimmomatic to trim the first 15 and last 45 bases and perform TruSeq3-PE-2 adapter trimming with parameters: *LEADING:3 TRAILING:3 SLIDINGWINDOW:4:15*. Reads were then mapped and processed with HISAT2 and Samtools using default quality settings. Read counting was performed similarly to single end reads and a single counts file was generated by merging the counts files for the single and paired end samples.

R (v.4.5.2) with tidyverse (v.2.0.0) and DESeq2 (v.1.50.2) (8) was used for all principal component and differential expression analysis. Since our samples from the single end (Spont, ZD, and GF) and paired end (Cycling and Starvation) RNA-sequencing data were collected separately and our experimental model including all the samples lacked a mixed-model approach, batch effect correction was not performed for DESeq2 analysis. Since the nature of the sequencing data violates the DESeq2 assumption that the majority of genes don’t change, all genes with differential expression (padj < 0.05) in binary comparisons were compiled and excluded from the final data set matrix and normalization was performed using the controlGenes parameter. Then DESeq2 was performed on the filtered normalized dataset. To get lists of differentially expressed genes, pairwise comparisons were performed between quiescence triggers(T) versus spontaneous quiescence(T/Spont) or triggers vs. Cycling(T/Cycling) conditions. For any pairwise comparisons, a p-adjusted(padj) value of < 0.05 for a particular gene was considered as a significant differentially expressed (DE) gene, whereas genes with padj > 0.05 were labelled as non-significant. Normalized counts from RNA-Seq data were Z-score transformed per gene across samples (mean subtracted, divided by standard deviation) prior to visualization. Heatmaps were generated by ComplexHeatmap(9) package (v. 2.26.0) in R with hierarchical clustering applied to genes and samples using Euclidean distance and linkage parameters. Heatmap in **Fig. 1C** was generated using a representative set of 8269 genes (padj< 0.05) identified from Starv/Cycling comparison (dendrograms not shown). Plots and analysis were carried out in R and GraphPad Prism 11, and Venn diagrams were generated using the web-based DeepVenn platform.(10)

**Analyzing metal associated transcriptomic changes**

To assess changes in metal-related gene expression, comprehensive gene lists for “metal ion homeostasis”, “metal ion dependent” and “metal ion transport” annotations were compiled from the Uniprot database for human entries found in Gene Ontology (GO). All reviewed entries with above annotations were downloaded and duplicates were removed prior to comparing with our RNA sequencing data. This resulted in a list of 2458 genes (**Dataset S2)** which was then used to find the significant differentially expressed genes (padj < 0.05) from T/Spont comparisons as shown in **Fig. 4*A*** and **Fig. S5**. It is worth noting that genes annotated under ‘Metal ion binding’ were excluded in this initial list. Metal ion binding genes were subsequently incorporated for a broader analysis, as described below. To analyze metal specific transcriptomic changes as presented in **Fig. 4*C-D***, separate lists for keyword annotations of “zinc ion”, “Fe ion”, “Cu ion”, “Mn ion”, “copper ion homeostasis,” “copper ion transport,” “manganese ion homeostasis”, and “manganese ion transport” for humans were downloaded from Gene Ontology with Uniprot database as a source for gene entries. The list of genes for the above annotations from the Gene Ontology were further curated to establish combined list of genes primarily associated with Cu and Mn ion “homeostasis and transport” by deleting the duplicate genes from each gene list for a specific metal ion. The curated list of genes associated with Cu and Mn ion “homeostasis and transport” has around 57 and 39 genes respectively (**Dataset S3, Fig. 4C)**. To get a broader picture of metal ion related genes, we combined the metal ion “binding”, “homeostasis”, “transport” related genes for each specific metal ion such as “zinc ion”, “Fe ion”, “Cu ion”, “Mn ion”, removed duplicates as described above and obtained a list with 2240 ‘zinc ion’ genes, 338 ‘Fe ion’ genes, 108 ‘Cu ion’ genes, and 94 ‘Mn ion’ genes (**Dataset S3,** **Fig.S*7A*)**. Additionally, Log2FC values of differentially expressed (padj< 0.05) Fe regulating genes (**Table 2**) corresponding to intracellular iron homeostasis, FeS cluster biogenesis, heme biosynthesis enzymes, and heme metabolism was plotted in **Fig. 5*A***. A curated list of 139 genes (**Dataset S4)** was further generated by combining associated genes from GO BP pathways for ‘heme metabolic processes’, ‘Intracellular Fe homeostasis’, and ‘Fe-S cluster assembly’ to subset DEGs (padj< 0.05) from T/Spont comparisons as presented in **Fig. 5*B***.

To identify significant DE genes involved in GOBP oxidative phosphorylation pathway and GOMF antioxidant activity pathway for T/Spont, that are also metal associated, we screened DEGs (padj<0.05) for genes annotated with ‘metal binding’, ‘metal dependent’, ‘metal homeostasis’ and any genes influenced by metal mediated stress which is reflected in **Fig. 6**. For each DEG a rank score was calculated as *rnk = -log10($padj) * sign($log2FoldChange)).* To examine the inverse relationship among quiescence conditions, DEGs were ranked using this score and plotted as shown in **Fig. S*7***. To identify the inverse relationship for ‘lysosome’ specific DEGs between ZD and Starv, a list was downloaded from GO for keyword annotations of “lysosome” with Uniprot database as a source for gene entries and ranked DEGs in Starv and ZD were plotted in **Fig. S7C**. Heatmap for metal associated genes in Spont, GF, ZD and Starv shown in **Fig. S5B** was generated using a representative set of 1221 metal associated DEGs (padj< 0.05) identified from ZD/Spont comparisons.

**Comparison of our dataset with publicly sourced data**

Our RNA-Seq experiment processed serum starvation induced quiescent samples separately from ZD-induced quiescence, GF withdrawal-induced quiescence, and spontaneously quiescent cells. As our analysis focused on differential gene expression among quiescence triggers particularly relative to spontaneous quiescence (Spont) and was performed without batch corrections. We validated the serum starvation dataset with a published dataset. Briefly, we compared our data with the publicly available GEO dataset (GSM3488784) from Min *et. al.*.(1) which includes RNA-seq profiles of sorted high p21-geminine MCF10A cells undergoing spontaneous quiescence in minimal media for 48h or 48h in serum starvation. Their experimental conditions closely matched with ours differing only by the addition of 0.3% Bovine serum albumin (BSA) in serum starvation. We performed DESeq2 analysis on the downloaded read counts dataset (GSM3488784) and performed a binary comparison between serum starvation and spontaneous quiescence conditions using our analysis pipeline to identify significant differentially expressed genes (DEGs, padj < 0.05). We found that 5,660 genes were significantly DE in Starv/Spont in Min *et. al.*’s dataset. To evaluate consistency with our data, we compared these DEGs with those identified in our dataset for Starv/Spont condition. We found that 4,132 genes out of 5,660 DEGs in Min *et. al.* dataset overlapped with ours (**Dataset S1 *and* Fig. S2D)**. The remaining differences likely come from the addition of BSA in their samples, as we know that albumin can bind to transition metal ions.(11) We further validated our results by examining “metal associated genes”. Notably, 80% of the metal associated DEGs in their dataset (570 out of 712 genes, padj < 0.05, **Dataset S2**) overlapped with our metal associated DEGs (padj < 0.05). Moreover, when we plotted the log2Foldchange of genes particularly involved in maintaining metal homeostasis, we observed that the downregulation and upregulation trends of those genes closely matched those observed in our analysis. Together, these analyses support the reliability of our RNA-seq results (**Fig. S2D** and **Fig. S5D**).

**Gene Ontology (GO) and Kyoto Encyclopedia of Genes and Genomes (KEGG) pathway analysis**

An online available pathway analysis tool “g:Profiler”(12) was used to identify Canonical Pathways, GO_Biological Processes, GO_Molecular Functions, GO_Cellular Component, KEGG analysis, and Reactome analysis. To perform the functional enrichment analysis from DE genes, the positively regulated DEGs and negatively regulated DEGs either with padj <0.05 or padj <0.01 values from each trigger/spontaneous comparisons were run separately for better understanding on top significant pathways in each quiescence conditions.

Figures

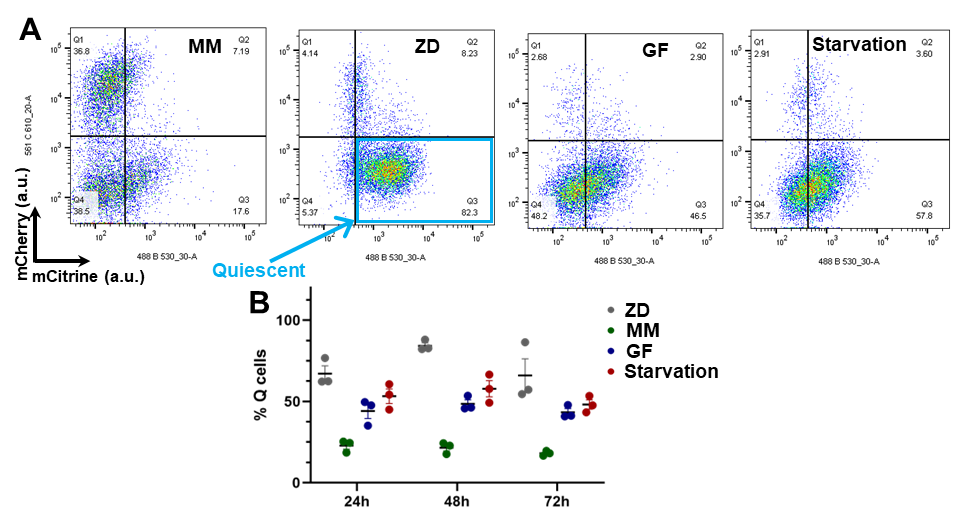

**Fig. S1.** **FACS analysis of quiescence in MCF10A p21-mCitrine geminin-mCherry cells**. (A) Representative scatter plots of mCherry vs mCitrine signals in cells grown for 48h in minimal (MM) media, ZD, GF, and Starv media. Each dot represents a single cell. Q3 quadrant represents MCF10A cells expressing high p21-mCitrine and low geminin-mCherry which identifies the quiescent population in the heterogeneous population of cells (cyan box). (B) Percentage of cells in the Q3 quadrant i.e. quiescent cells grown in ZD, GF, Starv and in minimal (Spont) media for 24h, 48 h, and 72 h. Each experiment was repeated in triplicate (N=3 biological replicates). Error bars represent SD.

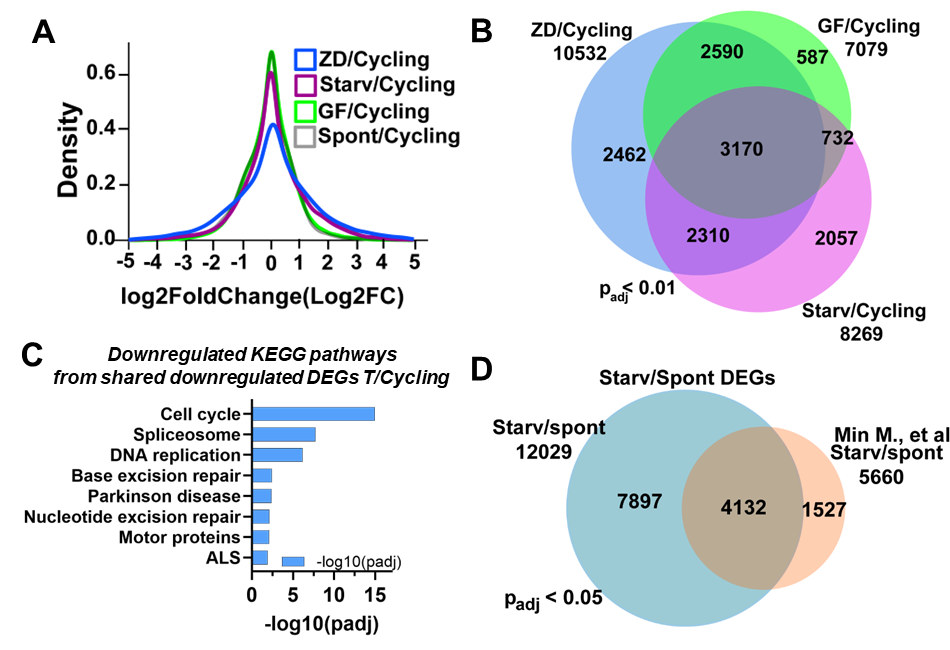

**Fig. S2. Transcriptomic analysis for quiescence triggers versus cycling conditions.** (A) Density plot showing Log2Foldchange (Log2FC) distributions of differentially expressed genes across all pairwise comparisons, Spont/Cycling, GF/Cycling, ZD/Cycling, and Starv/Cycling. (B) Venn diagram depicting differentially expressed genes (padj < 0.01) from analyzed binary comparisons of specific quiescence triggers vs cycling cells (ZD/Cycling, GF/Cycling, and Starv/Cycling). (C) KEGG pathway analysis highlighting key pathways enriched among the 1,230 commonly downregulated DEGs (padj < 0.01) across quiescence triggers/Cycling comparisons. (D) Comparative Venn diagram analysis of Starvation/Spontaneous quiescence sub-types from this study and reanalyzed Starv/Spont quiescence dataset from Min et al., *Plos Biology* (2019)(1). We found that 73% (4132/5660) of the DEGs reported by Wei et al overlap with our significant DEGs genes in Starv/Spont comparisons, with minor differences likely due to their addition of 0.3% bovine serum albumin (BSA) in the serum starvation cell culture media. Overall, this confirms the reliability of our analysis with trigger/spontaneous analysis (see methods).

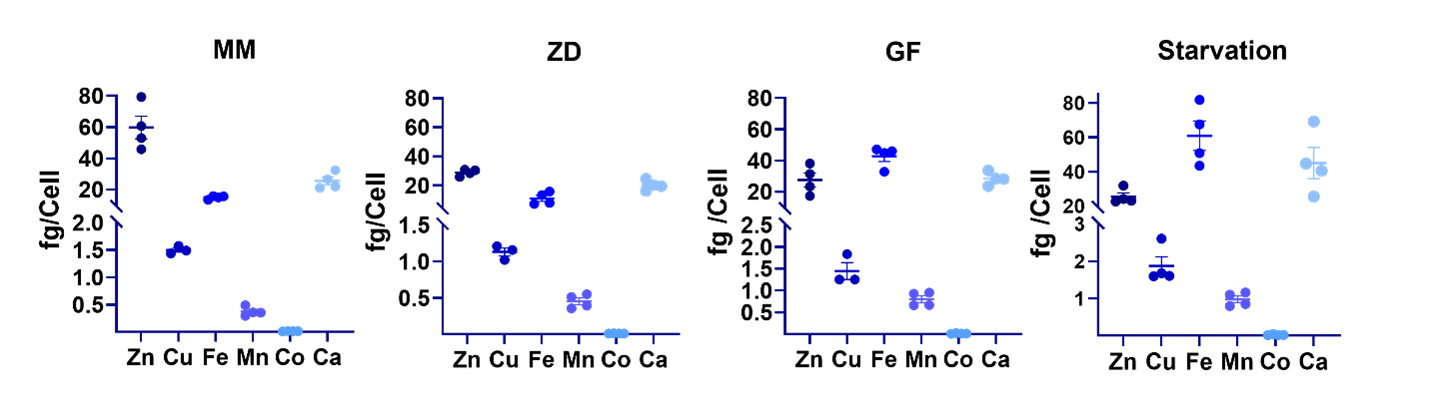

**Fig. S3.** **ICP-MS analysis of total amount of Zn, Cu, Fe, Mn, Ca, and Co in MCF10A p21-mCitrine geminin-mCherry** **cells** grown in MM (Minimal media), ZD (MM + 3 µM TPA), GF (MM without growth factor and insulin); Starv (MM without serum, growth factor and insulin) conditions for 48h. Each dot represents one biological replicate. The amount (fg) of metal was normalized to the cell count for each sample. Analysis was performed in N ≥ 3 biological replicates. Error bars represent SD.

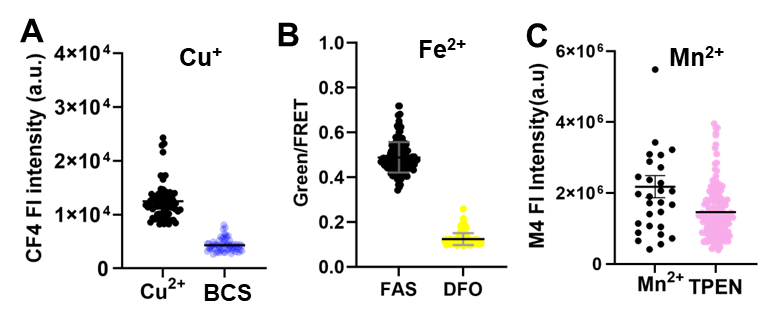

**Fig. S4. Live cell imaging with (A) Cu^+^, (B) Fe^2+^, and (C) Mn^2+^ ion specific probes with increase and decrease in metal content**. Increased metal condition: CuCl_2_ (100 µM, 1h), FAS (1 mM,1.5h), or MnCl_2_ (25 µM, 1h); Decreased metal condition: chelators BCS (500 µM,1 h), DFO (250 µM, 4 h), or TPEN (40 µM, 40 min). Cells treated with respective salts or chelators were washed with buffers, then treated with metal ion-specific fluorescence probes: CF4 (4 µM, 20 min, 37 ^o^C), FIP-1 (5 µM, 1.5 h, 37 ^o^C), M4 (5 µM, 15 min, 37 ^o^C) probes, followed by washing and imaging. Each dot represents the fluorescence intensity of the probe or intensity ratio coming from a single cell. Error bars represent SD.

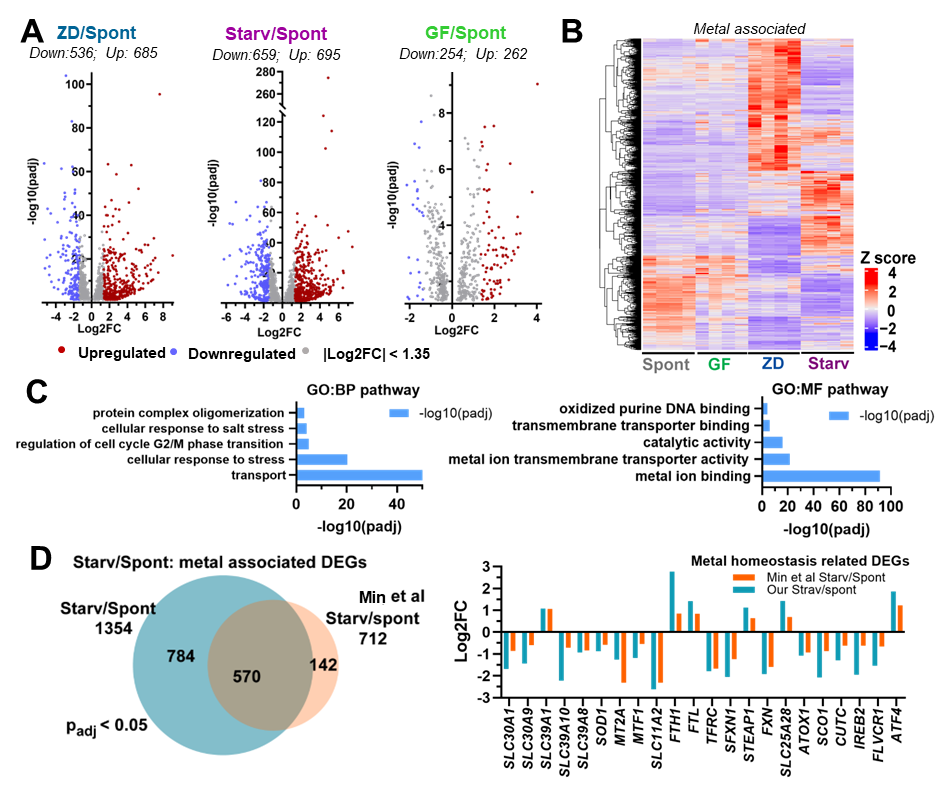

**Fig. S5. Metal associated DE genes and corresponding common pathways in quiescence triggers.** (A) Volcano plots of differential expression of genes (*p*adj < 0.05) in analyzed data groups. Each dot is a significant DE gene. Blue dots represent downregulated genes, and red dots represent up-regulated genes with Log2FC > |1.35|. Grey dots denote DEGs with Log2FC < |1.35|. (B) Heatmap displaying the expression level of metal associated differentially expressed genes (~ 1200 genes) in quiescence triggers. Relative gene expression levels are shown as Z-scores and color-coded from violet (low expression) to red (high expression), with white indicating zero. (C) Top five significant Gene ontology biological process (BP) and molecular function (MF) pathways enriched among 337 common metal associated DEGs (padj < 0.05) in specific quiescence triggers. (D) Venn diagram comparing metal associated DEGs in Starvation/Spontaneous from this study with the reanalyzed dataset from Min et al 2019 (1). Accompanying plot shows Log2FC expression of DEGs in a curated metal homeostasis gene set across both datasets.

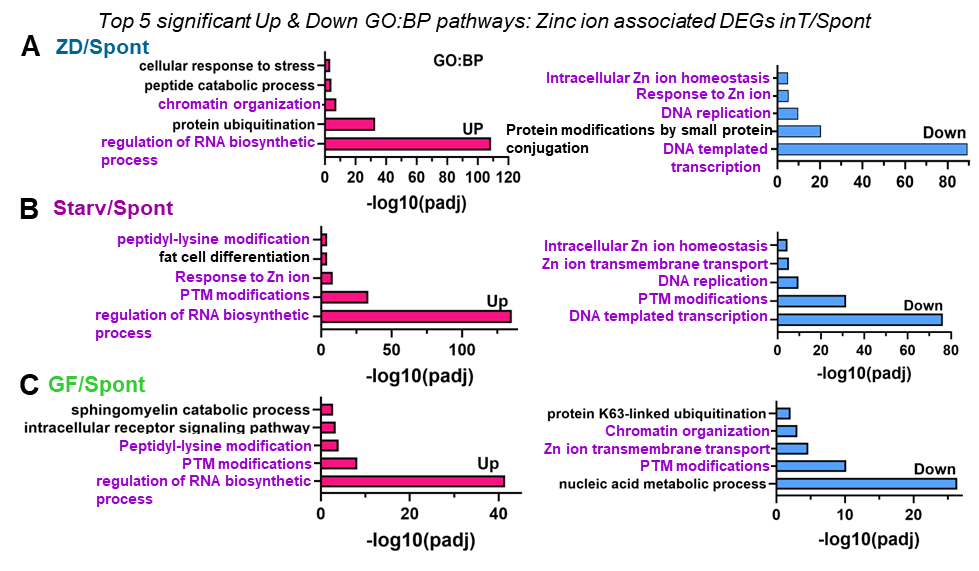

**Fig. S6. Significantly enriched zinc associated Gene ontology biological process pathways in T/Spont.** Top five significant upregulated and downregulated GO: BP pathways enriched from zinc associated DEGs (padj < 0.05) in ZD/Spont (A), Starv/Spont (B) and GF/Spont (C). Purple denotes the shared pathways in different quiescence triggers.

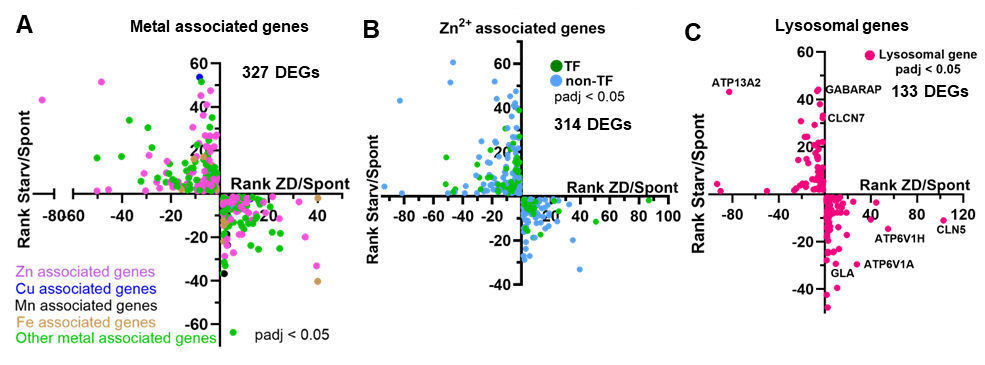

**Fig. S7.** **Reciprocal regulation relationship of DEGs between ZD/Spont and Starv/Spont.** (A) Plot showing ranked metal associated DEGs (padj < 0.05) inversely regulated in ZD/Spont and Starv/Spont, color coded specifically for zinc (purple), copper(blue), manganese(black), iron(brown) and other metal association(green). (B) Plot of oppositely regulated 314 ranked Zinc (Zn^2+^) associated DEGs (padj < 0.05) in ZD/Spont and Starv/Spont color coded specifically for transcription factor (TF, deep green) and non-TF (cyan). (C) Plot of 133 lysosomal genes (padj< 0.05) inversely regulated in ZD/Spont and Starv/Spont.

**Tables**

**Table S1. Imaging parameters for labile metal ion imaging in live cells.**

| **Cell type** | **Conditions** | **M^n+^ salt**  **(conc, time)** | **Chelator**  **(conc, time)** | **Probe**  **(conc, time)** | **Nuclear dye**  **(conc, time)** | **λ_excitation ,_ nm (% power, exposure time)** | **λ_emission,_ nm** |
| --- | --- | --- | --- | --- | --- | --- | --- |
| **Labile Cu^+^ imaging** | | | | | | | |
| MCF10A  wildtype | Positive control | CuCl_2_  (100 µM, 1h) | n.a. | CF4  (4 µM, 20 min, 37 ^o^C) | n.a | 20X air objective, NA 0.75  514 nm  (30, 300 ms)  Dichroic ZT445/514/594rpc | 525–555 |
|  | Negative control | n.a | BCS  (500 µM,1 h) |  | n.a |  |  |
|  | 3ZD or MM or GF or Starv | n.a | n.a |  | n.a |  |  |
| **Labile Fe^2+^ imaging** | | | | | | | |
| MCF10A H2B-Halo | Positive control | Ferrous ammonium sulphate FAS  (1mM,1.5 h)  freshly prepared in degassed water | n.a. | FIP-1  (5 µM, 1.5 h, 37 ^o^C) | JF669  (10 nM, 15 min, 37 ^o^C) | 20X air objective, NA 0.75  Green:488 (40, 300 ms)  FRET:488 (40, 300 ms)  Red: 561 (10, 300 ms)  Cy5_ex_: 640 (10, 300 ms)  Dichroic ZT405/488/561/640rpcv2, ZT445/514/594rpc | Green:  GFP_em_  (500-550)  FRET: TRITC_em_ (575-625)  Red: TRITC_em_ (575-625*)*  Cy5_em_: 671-739 |
|  | Negative control | n.a | DFO  (250 µM,4 h) |  |  |  |  |
|  | 3ZD or MM or GF or Starv | n.a | n.a |  |  |  |  |
| **Labile Mn^2+^ imaging** | | | | | | | |
| MCF10A  wildtype | Positive control | MnCl_2_  (25 µM, 1h) freshly prepared in water | n.a. | M4  (5 µM, 15 min, 37 ^o^C) | n.a. | 20X air objective, NA 0.75  488  (80, 600 ms)  DichroicZT405/488/561/640rpcv2, | 500-550 |
|  | Negative  control | n.a | TPEN  (40 µM,40 min) |  | n.a |  |  |
|  | 3ZD or MM or GF or Starv | n.a | n.a |  | n.a |  |  |
| **Labile Zn^2+^ imaging** | | | | | | | |
| MCF10A  NES-ZapCV2 H2B-Halo | 3ZD or MM or GF or Starv | n.a | n.a | Genetically encoded cytosolic ZapCV2 sensor | JF669  (10 nM, 15 min, 37 ^o^C) | 10X or 20X air objectives  CFP:440  (50, 400 ms) 455 dichroic  FRET:440 (50, 400 ms)  455 dichroic  Cy5 _:_640  (50, 200 ms)  640 dichroic | FRET: YFP_em_ 519-561  CFP_em_  460-500,  Cy5_em_:683-727 |

**Table S2. Iron regulating genes in ZD/Spont and Starv/Spont.**

|  | **ZD / Spont** | **Starv / Spont** |
| --- | --- | --- |
| *Fe uptake* | **Upregulated**  ***TFRC*** – Fe uptake  ***CYBRD*1** – ferric reductase  ***STEAP4*** – ferrireductase enabling fe import  ***SCARA5*** – ferritin receptor  ***SLCO2B1*** – heme/porphyrin transporter | **Downregulated**  ***TFRC*** – major transferrin-mediated iron uptake receptor  ***IREB2*** – iron starvation regulator |
| *FeS biogenesis* | **Downregulated**  ***BOLA3*** – FeS cluster maturation factor  ***GLRX5*** – transfers FeS clusters to recipient proteins  ***FXN*** – frataxin, provides iron for cluster assembly  ***MMS19*** – cytosolic FeS protein maturation  ***SLC25A39*** – mitochondrial glutathione import required for FeS metabolism  ***ABCB7*** – exports FeS cluster precursors from mitochondria to cytosol | **Downregulated**  ***BOLA3*** – FeS cluster maturation factor  ***GLRX5*** – transfers FeS clusters to recipient proteins  ***FXN*** – frataxin, provides iron for cluster assembly  ***NDUFAB1*** – essential subunit of Complex I and coordinates assembly of other complexes (stabilizes FeS biogenesis complex) |
| *Heme metabolism* | **Dysregulation**  ***ALAS1*** – rate limiting enzyme in heme synthesis **increases**  ***HMOX1*** – heme degradation **increases**  ***ABCB6*** –porphyrin transport **increases**  ***BLVRB*** – biliverdin reductase **increases**  ***CPOX*** – sixth enzyme in heme biosynthetic pathway **decreases**  ***HMBS*** – third enzyme in heme biosynthesis pathway **decreases** | **Dysregulated**  ***HMOX*** – heme degradation **increases**  ***PPOX*** – seventh enzyme in heme biosynthetic pathway **increases**  ***BLVRB*** – biliverdin reductase **increases**  ***HPX*** – heme sequestering **increases**  ***HMBS*** – third enzyme in heme biosynthesis pathway **decreases**  ***UROS* –** fourth enzyme in heme biosynthetic pathway **decreases**  ***CPOX*** – sixth enzyme in heme biosynthetic pathway **decreases** |
| *Signature of high Fe* |  | ***LCN2*** – binds iron chelating compounds to regulate iron availability **increases in high Fe**  ***HIF1A*** – hypoxia induced TF is **downregulated in high Fe**  ***ATP6V1A*** – lysosomal acidification and activation of Fe-dependent prolyl hydroxylases **decreases in high Fe**  ***ATP6V01B* or *ATP6V0A2*** – lysosomal acidification and activation of Fe-dependent prolyl hydroxylases **decreases in high Fe** |
| *Lysosomal and Fe handling pathways* | **Downregulated**  ***ATP13A2*** – lysosomal transport protein important for polyamines, metal transport, and mitophagy – decreases | **Upregulated**  ***ATP13A2*** – lysosomal transport protein important for polyamines, metal transport, and mitophagy  ***ATP6AP1*** – lysosome acidification  ***CCDC115*** – Golgi homeostasis, intracellular trafficking, iron and heme metabolism |
| *Fe sequestration and storage* | **Upregulated**  ***FTH1*** – ferritin heavy chain  ***FTL*** – ferritin light chain | **Upregulated**  ***fth1*** – ferritin heavy chain |
| *Fe transport* | ***SLC25A28* –** mitochondrial membrane Fe importer, essential for mitochondrial function and FeS biogenesis, **decreases**  *Fe export increases*  ***SLC40A1*-** plasma membrane Fe exporter, **increases**  ***FLVCR1*-** responsible for heme export, **decreases** | ***SLC25A28, SLC25A37* –** mitochondrial membrane Fe importer, essential for mitochondrial function and FeS biogenesis, **upregulated**  ***SLC11A2* –** plasma membrane Fe import, **upregulated**  ***FLVCR1*-** responsible for heme export, **decreases** |

**Dataset S1(separate file).** List of differentially expressed gene (padj < 0.01) for T/Spont and T/Cycling comparisons.

**Dataset S2 (separate file).** List of 2458 metal associated gene downloaded from Gene Ontology and list of “metal associated” DE genes (padj < 0.05) for the different triggers/Spont.

**Dataset S3 (separate file).** Comprehensive list of genes for ‘Zinc ion’, 'Cu ion', 'Mn ion', 'Fe ion', ‘Cu ion homeostasis and transport’, ‘Mn ion homeostasis and transport’ from Gene Ontology along with corresponding DEGs for zinc ion as well and Cu/Mn homeostasis and transport across each T/Spont comparisons.

**Dataset S4 (separate file).** Comprehensive list of genes for iron homeostasis.
